# Restored alveolar epithelial differentiation and reversed human lung fibrosis upon Notch inhibition

**DOI:** 10.1101/580498

**Authors:** RM Wasnick, M Korfei, K Piskulak, I Henneke, J Wilhelm, P Mahavadi, D von der Beck, M Koch, I Shalashova, O Klymenko, L Fink, H Witt, H Hackstein, E El Agha, S Bellusci, W Klepetko, M Königshoff, O Eickelberg, T Braun, W Seeger, C Ruppert, A Guenther

**Affiliations:** Department of Internal Medicine, Justus-Liebig-University Giessen, 35392 Giessen, Germany; Institute of Pathology, UEGP, 35578 Wetzlar, Germany; Department of Pediatrics, Technical University Munich, 85354 Freising, Germany; Department of Clinical Immunology and Transfusion Medicine, 35392 Giessen, Germany; Department of Thoracic Surgery, Vienna General Hospital, 1090 Vienna, Austria; Comprehensive Pneumology Center, Research Unit Lung Repair and Regeneration, Helmholtz Center Munich, Ludwig Maximilians University Munich; 81377 Munich, Germany; Max-Planck-Institute for Heart and Lung Research, 61231 Bad Nauheim, Germany; Lung Clinic Waldhof-Elgershausen, 35753 Greifenstein, Germany; Excellence Cluster Cardio-Pulmonary System (ECCPS), 35392 Giessen, Germany; Cardio-Pulmonary Institute (CPI), 35392 Giessen, Germany; University of Giessen and Marburg Lung Center (UGMLC); German Center for Lung Research (DZL), 35392 Giessen, Germany; European IPF Registry / Biobank, 35392 Giessen, Germany; Division of Respiratory Sciences and Critical Care Medicine, University of Colorado, United States; Department of Medicine, Division of Pulmonary Sciences and Critical Care Medicine, University of Colorado, United States

**Author notes:** To whom correspondence should be addressed: Andreas Günther, MD, Prof of Internal Medicine, Director “Centre for Interstitial and Rare Lung Diseases”, University of Giessen Medical School, Justus-Liebig-University Giessen, Klinikstrasse 36, 35392 D-Giessen, +49 641/98542502. both authors contributed to the manuscript equally.

**Keywords:** Lung fibrosis, ILD, DPLD, Notch, lung surfactant, alveolar type 2 cell, alveolar surface tension, aspartyl protease, epithelial injury, epithelial regeneration

## Abstract

Alveolar epithelial cell type II (AEC2) injury underlies idiopathic pulmonary fibrosis (IPF). Here we show increased Notch1 signaling in AEC2s in human IPF and IPF models, causing enhanced proliferation and de-differentiation of AEC2s. As a result, we observed defective surfactant protein (SP)-B/C processing, elevated alveolar surface tension, repetitive alveolar collapse and development of lung fibrosis. Similar changes were encountered upon pharmacological inhibition of SP-B/C processing in vivo by pepstatin A. Inhibition of Notch signaling in cultured human IPF precision cut lung slices improved surfactant processing capacity of AEC2s and reversed fibrosis. Notch1 therefore offers as novel therapeutic target.

**One sentence summary:** Notch1 inhibition restores alveolar epithelial differentiation and surface tension and reverses matrix deposition in lung fibrosis

## Introduction

Idiopathic pulmonary fibrosis (IPF) is a progressive, devastating, and ultimately fatal lung disease ^1^. The pathomechanism of IPF is not fully understood, but it appears to be multifactorial in nature, involving a predisposing genetic background as well as exogenous second hits ^2 3 4^. Chronic alveolar epithelial type 2 (AEC2) injury plays a pivotal role in IPF ^5–7^, and the observed epithelial abnormalities include hyperplastic as well as apoptotic AEC2s in close proximity to the fibroblast foci, and cystic airspaces (microscopic honeycombing) covered by either alveolar or bronchiolar-like epithelium ^8^.

One central function of AEC2s is to produce, process and adequately secrete pulmonary surfactant, a complex lipoprotein mixture that covers the alveolar space and lowers alveolar surface tension to less than 5mN/m instead of ~70mN/m, as in case of pure water, making breathing possible at normal transpulmonary pressure gradients ^9^. In addition to a relatively high amount of dipalmitoylated phosphatidylcholine, pulmonary surfactant contains the two extremely hydrophobic proteins SP-B and SP-C, both of which are of pivotal importance for lowering the alveolar surface tension. Because of their fusogenic properties (they easily attack cell membranes in their mature forms), SP-B and SP-C are synthesized as pro-proteins (42 and 21 kDa, respectively), which undergo sequential proteolytic cleavage by aspartyl and cysteine proteases including napsin A (SP-B), cathepsin H (SP-B/C), and pepsinogen C (SP-C) during their journey through the lysosomal compartment in AEC2s (trans Golgi, multivesicular bodies, lamellar bodies) ^10^.

Another key function of AEC2 is to act as progenitors for the alveolar epithelium, as they are able to self-renew and differentiate into alveolar epithelial type 1 cells (AEC1s) in response to alveolar epithelial damage ^11,12^. Of interest, reactivation of developmental pathways in AEC2 like sonic hedgehog ^13^, TGFβ ^14^, and Wnt ^15^ has been previously demonstrated in IPF. Despite evidence suggestive of altered regulation of Notch signaling in epithelial cells of IPF lung tissues, the roles for Notch in disease pathogenesis and altered epithelial cell function are poorly defined ^16^. Notch signaling relies on four receptors (NOTCH1 through NOTCH4) that bind to and are activated by five ligands (DLL1, 3, and 4 and Jagged1 and 2) ^17^. Upon ligand binding, the Notch receptor undergoes two cleavage events mediated by the γ**-**Secretase and Presenilin complex. This results in the release of the Notch intracellular domain (NICD), which translocates into the nucleus and activates transcription of downstream target genes ^17,17^. In the lung, *Hes1* is the best-characterized Notch1 downstream target gene ^18^. Notch plays a major role as a mediator of cell-cell communication by modulating cell proliferation and differentiation ^19^.

Based on an initial transcriptome analysis of IPF and donor lung tissues, which revealed differential regulation of Notch signaling in IPF, we hypothesized that this pathway plays an important role in AEC2 proliferation and sought to determine the consequences of Notch activation on AEC2 differentiation. In this study we demonstrate that, in IPF and the bleomycin model of lung fibrosis, increased Notch1 signaling in AEC2s induces their proliferation and de-differentiation, resulting in defective surfactant protein (SP)-B/C processing and elevated alveolar surface tension ex vivo, in vitro and in vivo.

## Results

### Expiratory alveolar collapse in patients with IPF

Increased intra-pulmonary shunt-flow, suggestive of alveolar collapse, has been proven in IPF patients in the past ^20^.To provide further evidence for subpleural, repetitive alveolar collapse during the respiratory cycle of IPF patients, we used high-resolution (HR) CT scans obtained during inspiration and end-expiration from IPF and COPD patients and from patients who were not suffering from structural lung diseases (controls, information on patient cohorts in Table S1). In a first approach, hand-segmented transversal CT layers were color-coded for density: in IPF subjects there was an increase in subpleural density during expiration, corresponding to an increase in non-aerated lung areas (example shown in Fig. 1A). Next, we analyzed statistically the density distribution in the basal layers of IPF, COPD and control patients in inspiration and expiration in a pooled fashion. Fig. 1B, left panel, displays the average densitogram over the entire Houndsfield Unit (HU) spectrum from non-aerated, poorly aerated, normally aerated to hyperinflated lung areas ^21,22^. As evident from this graph, IPF patients showed a higher percentage in lung areas being poorly or non-aerated and this fraction appeared highly increased at the end of expiration. Likewise, when analyzing these data in a categorial approach (Fig. 1B, right), we observed a significantly increased percentage of lung areas being poorly or non-aerated in IPF (n=24) vs COPD (n=5) or Control (n=4) at the end of expiration. Finally, the total subpleural space of hand-segmented transversal CT layers obtained at three different positions (apical, bifurcation, and basal) was analyzed for the HU distribution and shift during expiration. IPF patients showed a significantly higher increase in subpleural density at the end of expiration than COPD subjects and controls (Fig. 1C). Of note, this increase was not only visible in the lower parts of the IPF lungs, where the disease is usually more prominent, but also in the most apical layers, where fibrotic changes are usually less apparent ^23^. Taken together these data clearly demonstrate an increase in alveolar collapse in IPF patients, most likely leading to extensive and repetitive alveolar epithelial cell injury.

**Fig. 1.**
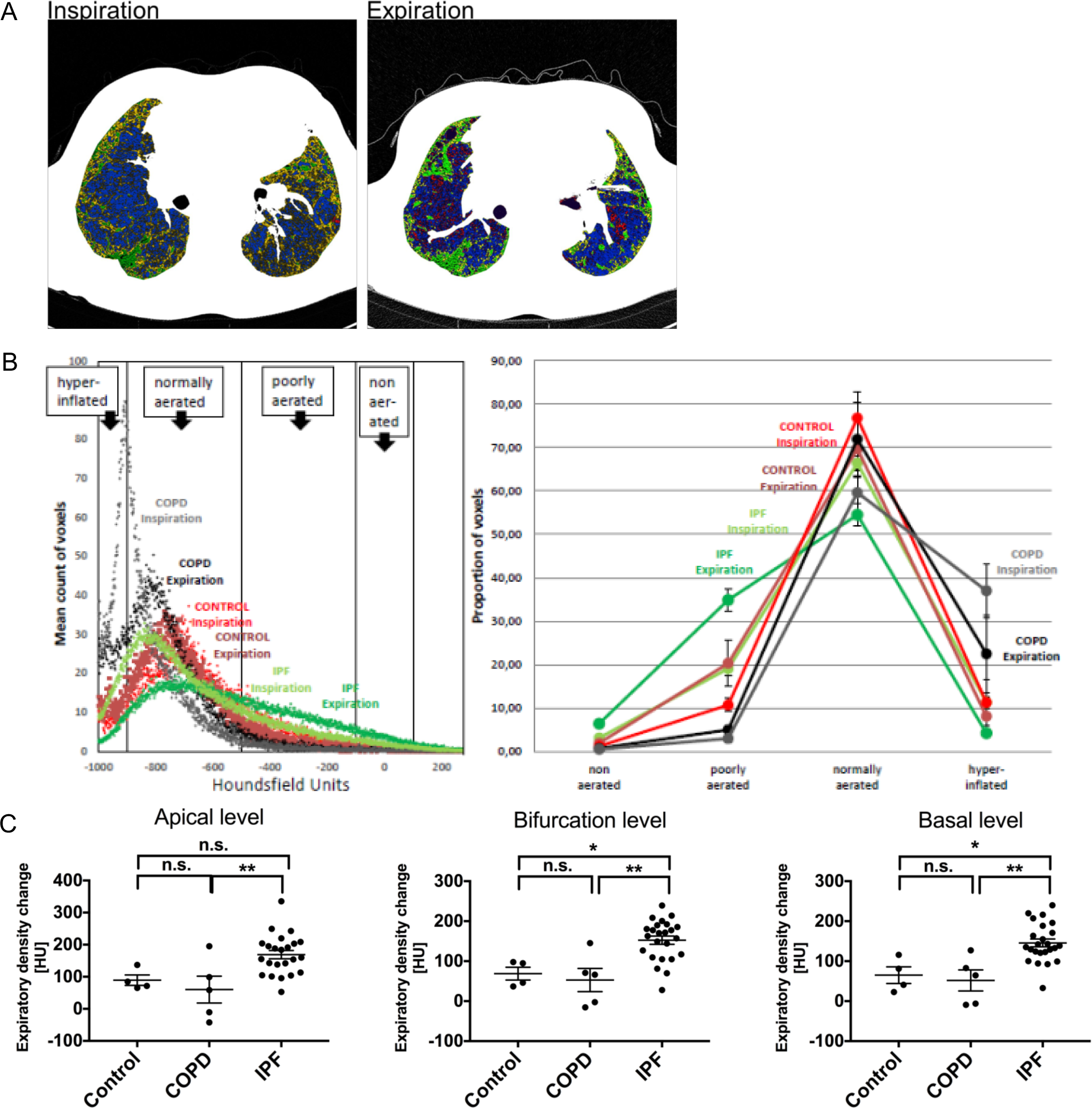
Expiratory alveolar collapse in patients with IPF. (A) Representative HRCT from one IPF patient during inspiration and expiration. Voxels were color-encoded according to their density in houndsfield units (HU: hyperinflated = – 1000 to −901 = red; normally aerated = −900 to −501 = blue; poorly aerated = −500 to −101 = yellow; non-aerated = HU −100 to 100 = green). Note the preferentially subpleural increase in green areas in expiration indicating alveolar collapse. (B) Density distribution in HU in basal horizontal HRCT sections from pooled IPF (n=24), COPD (n=5) and Controls (n=4) during inspiration and expiration (left) and categorical distribution of voxels into non-aerated, poorly aerated and normally aerated as well as hyperinflated lung areas (right). Individual sections were hand-selected, transferred and analyzed for HU distribution. All data from all patients per group were pooled and are given as mean densitogram for the 3 patient categories and inspiration versus expiration (left). Right panel shows percentage of voxels per each density group for the same patient groups in inspiration and expiration. For the non-aerated tissue, differences reached significance for IPF vs Control (p=0.027 at expiration; 0.057 at inspiration), for IPF versus COPD (p<0.001 at expiration; p=0.006 at inspiration), but not for Control versus COPD. In the poorly aerated tissue, differences reached significance for IPF vs Control (p=0.034 at expiration; p=0.032 at inspiration), for IPF versus COPD (p<0.001 at expiration; p<0.001 at inspiration), and for Control versus COPD (p=0.016 at expiration; p=0.016 at inspiration; all Wilcoxon-Mann-Whitney Test). (C) Quantification of density changes in corresponding subpleural HRCT sections at the basal, tracheal bifurcation, and apical level in expiration versus inspiration in the same patients as outlined in (D). Data are presented as the mean ± SEM. *p < 0.05, **p < 0.01, ***p < 0.001, n.s. = not significant by ANOVA

### Loss of Alveolar Surface Activity in IPF Is Primarily Based on Loss of Mature SP-B

The most probable cause of increased alveolar collapse in IPF is the increase in surface tension at the alveolar air-liquid interface ^24^, which is normally maintained at very low levels by the surfactant system. This prompted us to analyze its composition in the BALF of healthy volunteers and IPF subjects, as well as in lung homogenates from age-matched organ donors, COPD and IPF patients undergoing lung transplantation. In BALF, a prominent loss of mature hydrophobic surfactant proteins SP-B (Fig. 2A - BALF) and SP-C (Fig. S. 1A) was encountered in all IPF patients investigated. Moreover, the 23-kDa C-terminal proSP-B intermediate (CproSP-B, Fig. 2A) and the 21-kDa proSP-C (Fig. S. 1A) were detected in BALF from IPF patients (n=10 IPF patients, n=8 healthy volunteers). In explanted lung tissue from IPF patients, a similarly marked loss of mature SP-B (Fig. 2A and Figure S.1B - Tissue) and SP-C (Fig S. 1A) was evident and was accompanied by pronounced accumulation of the 42-kDa SP-B precursor, the 23-kDa C-terminal and the 20-kDa N-terminal proSP-B peptides, and smaller C-terminal processing intermediates of SP-B in the range of 12–16 kDa (Fig. 2A – Tissue, Fig. S. 1B; n=20 IPF patients, n=9 COPD patients and n=12 donors). As a result, we observed a massive reduction in the ratio of mature to total SP-B in IPF lung tissues (Fig. 2A). Moreover, this reduction was paralleled by a decrease in surfactant processing peptidases Napsin A and Cathepsin H (Fig. 2A and B, Fig. S. 1B; n=9 IPF patients, n=9 COPD patients, n=12 donors). Consequently, the amount of proSP-B and CproSP-B was significantly increased in the serum of IPF patients compared to donors, suggestive of increased epithelial barrier permeability, following epithelial injury and/or basolateral secretion (Fig. 2C; n=5 healthy controls, n=5 IPF). Tissue stores of phosphatidylcholine and phosphatidylglycerol, the two major surfactant phospholipids, were also significantly reduced in IPF (n=14) lung compared with lung tissue from the age-matched COPD (n=7) patients and from donors (n=9), also suggesting a loss of surfactant-specific differentiation of AEC2s (Fig. S. 1E). Genotyping of SP-B, SP-C and NAPSIN A, did not reveal any mutations in the investigated IPF population (Table S2), confirming that post-translational processing was altered in IPF patients.

**Fig. 2.**
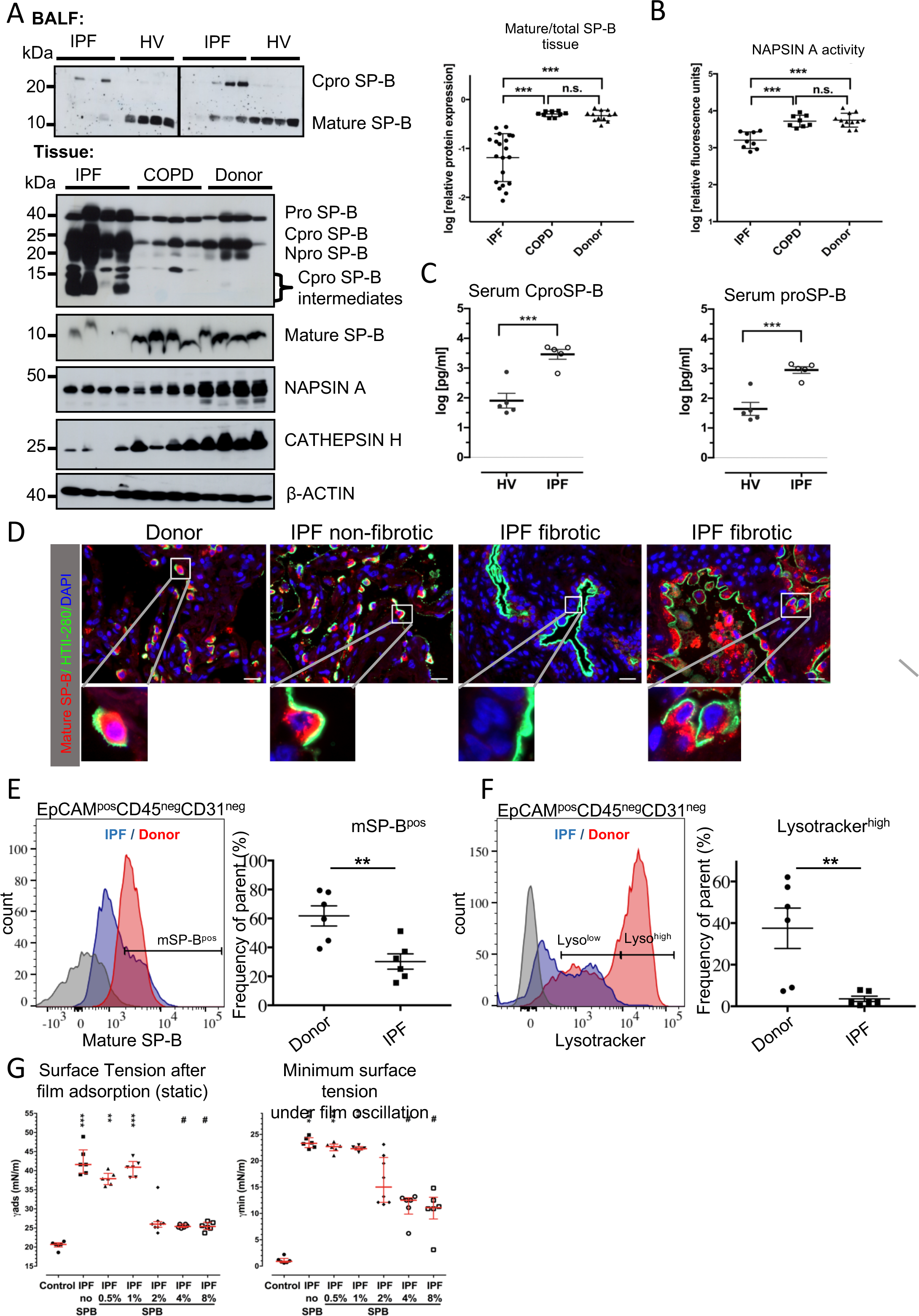
Loss of alveolar surface activity in IPF is primarily based on loss of mature SP-B (A)Upper panels: western blot analysis for mature SP-B in the BALF of IPF patients (n = 10) and healthy volunteers (n = 8). Lower panels: western blot analysis for mature SP-B, Napsin-A and Cathepsin-H in the peripheral lung tissue of IPF (n = 20), COPD (n = 9), and human donors (n = 12). The ratio of the mature to total SP-B in the peripheral lung tissue in shown in the adjacent graph. Data presented as mean values± SEM: ***p < 0.001, n.s. = not significant, by Student’s t-test. (B) NAPSIN A activity in peripheral lung tissue from patients with IPF (n = 9) and COPD (n = 9) and from human donors (HD, n = 12), as determined with a fluorogenic substrate. Data presented as mean values± SEM: ***p < 0.001, n.s. = not significant, by Student’s t-test. (C) Serum levels of proSP-B and CproSP-B in patients with IPF (n = 5) and healthy controls (n = 5). Data presented as mean values± SEM: ***p < 0.001 by Student’s t-test. (D) Representative immunofluorescence staining for HTII (green) and mature SP-B (red) in human donor and IPF lung tissue (n = 9 donors, n = 10 IPF patients). Scale bars, 20 µm. (E), (F) Representative histograms of flow cytometry analysis of mature SP-B (E) and Lysotracker incorporation (F) in the epithelial compartment of the lung defined as the EpCAM^pos^ CD45^neg^ CD31^neg^. The proportion of mature SP-B^high^ and Lysotracker^high^ were quantified in the adjacent graphs (n=6 donors, n=6 IPF patients). Data presented as mean values± SEM: **p < 0.01 by Student’s t-test. (G) Analysis of surface tension (□) in LSA pools from healthy volunteers (n=5) and IPF patients (n=15) at the end of 12sec of film adsorption (static – left graph) and under dynamic film oscillation in the Pulsating Bubble Surfactometer following mature SP-B supplementation (n=5 healthy volunteers and n=15 IPF patients). Given is the minimum surface tension after 5min film oscillation at a rate of 20 x/min (right graph). Data are presented as mean ± SEM: ** p < 0.01 and *** p < 0.001 when compared to controls and # p<0.05 when compared to IPF without SP-B by ANOVA

Using immunohistochemistry, mature SP-C (Fig. S. 1C), and mature SP-B (Fig. 2D and Fig. S. 1D), both of which co-localized with the AEC2 marker HTII-280 or ABCA3 in donor tissues (n=9), were expressed in a similar pattern in normal appearing areas of the fibrotic lungs (n=10). However, in the fibrotic areas, expression of mature SP-B and SP-C was heterogenous, with areas of adjacent, hyperplastic HTII-280^pos^AEC2s expressing similar levels of mature SP-B mixed with areas of HTII-280^pos^ AEC2s devoid of mature SP-B or mature SP-C (Fig. 2D and Fig. S. 1C, D). To quantify the size of the surfactant processing compartment at single cell level, we determined by flow cytometry the amount of intracellular mature SP-B and the capacity of the lamellar bodies of AEC2s (Lysotracker uptake -references ^25,26^) in the epithelial compartment, defined as EpCAM^pos^ CD45^neg^ CD31^neg^ Live population (Fig. 2E and 2F) from n=6 donor and n=6 IPF samples. IPF patients had a statistically significant decrease in the number of mature SP-B^pos^ epithelial cells (Fig. 2E), which was paralleled by a decrease in the in the number of Lysotracker^high^ epithelial cells (Fig. 2F), demonstrating that in IPF alveolar epithelial cells indeed have a lower capacity to process surfactant.

To determine if these changes in surfactant protein processing occurred earlier during the course of IPF, we also analyzed surgical biopsies from IPF patients obtained at the time of diagnosis and we observed similar disruptions in the processing of the hydrophobic SP as described above for tissue samples obtained at the time of lung transplantation (Fig. S. 1F, n=4 donors, n=4 IPF LTX explant tissue, and n=4 open lung biopsy). Collectively, these results indicate that altered SP processing is a consistent feature in fibrosis development seen in patients with IPF regardless of the stage of the disease.

When analyzing the surface activity of a pooled (n=15 patients) large surfactant aggregate (LSA) fraction of IPF BALF, we reconfirmed an extensive loss of surface activity in IPF versus healthy controls. Supplementation of this pooled IPF LSA with increasing doses of mature bovine SP-B (0.5% to 8%; wt/wt) alone was sufficient to greatly improve surface activity in a dose-dependent manner, reaching almost normal surface tension values (n=5 healthy volunteers) upon film adsorption (γads) and strikingly improved minimum surface tension (γmin) values at a physiological relative SP-B concentration of 2-4% (Fig. 2G) ^9^.

### Inhibition of SP-B Processing Recapitulates all Features of Lung Fibrosis

We next investigated the cellular consequences of the intracellular accumulation of proSP-B encountered in IPF. We reasoned that blocking SP processing would lead to intracellular accumulation of the pro- and intermediate forms, induction of ER stress, a prominent feature in IPF AEC2s, as well as AEC2 apoptosis and lung fibrosis. SiRNA-mediated downregulation of *napsin A* in MLE12 cells (Figure 3A, C resulted in the accumulation of 42-kDa proSP-B and proSP-C intermediates and induction of ER stress, as indicated by XBP-1 splicing and increased ATF6 expression (Fig. 3A, B).

**Fig. 3.**
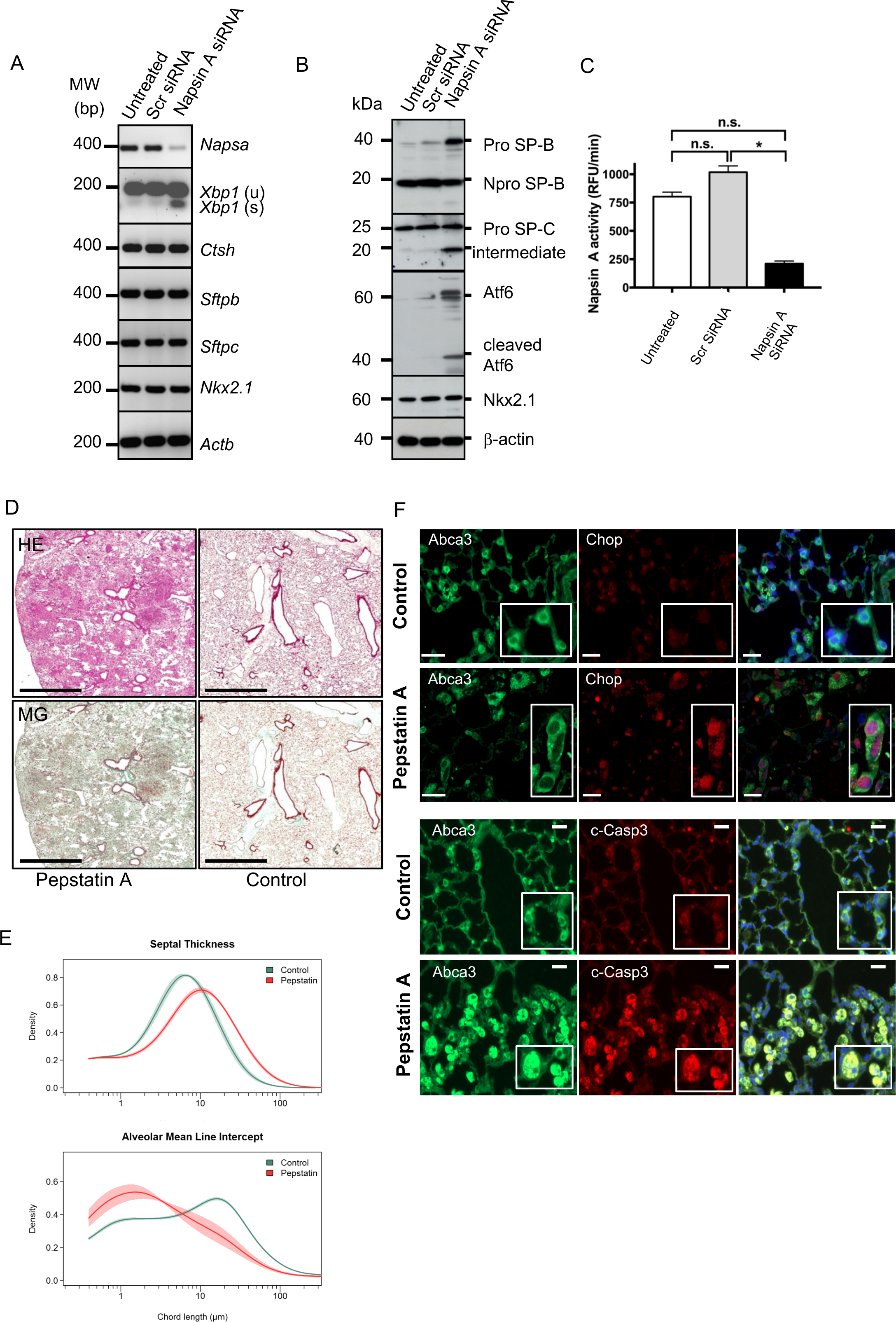
Inhibition of SP-B processing recapitulates all features of lung fibrosis. (A) RT-PCR analysis of *Napsa*, spliced (s) and unspliced (u) *Xbp1, Ctsh, Sftpb, Sftpc*, and *Nkx2.1* transcripts in MLE12 cells 2days after *Napsa* siRNA transfection. (B) Western blot analysis of proSP-B, proSP-C, Atf6 (ER-stress marker), and Nkx2.1 in *Napsa* siRNA– transfected MLE12 cells 6 days after transfection. Untreated cells and cells transfected with a scrambled siRNA served as the controls. (C) Napsin A activity in MLE12 cells 2 days after transfection with *Napsa* siRNA. Data presented as mean values± SEM: *p < 0.05; n.s.=not significant by Student’s t-test. (D) Representative images of low-field magnification of lung tissue sections stained with H&E (upper panel) and Masson’s trichrome (lower panels) from mice treated with pepstatin A for 28 days or controls. Scale bars, 1 mm. (E) Morphometric analysis of lung sections from mice described in (D). Morphometry data are shown as mean values of septal thickness and alveolar mean linear intercept from individual sections and as a probability density estimate function. The displayed curves represent point-wise mean values from individual densities within a group, and the shaded area indicates the standard errors. (F) Representative immunofluorescence staining for Abca3 (green) and either Chop or cleaved caspase-3 (red) of tissue sections from the mice described in (D). Scale bar = 50 µm in Chop/ABCA3 images, 20 µm in cleaved Caspase3/Abca3 images.

To inhibit SP-B processing *in vivo*, we administered the Napsin A inhibitor pepstatin A transbronchially into the lungs of C57BL/6N mice daily for 28 days, which resulted in a marked increase in proSP-B, but not proSP-C in the lungs of treated animals (Fig. S. 2A, B). Pepstatin A–treated mice (n=4 treated, n=4 untreated) showed notably reduced survival (pepstatin A: 70%; control: 100%) and a significant loss of pulmonary compliance (4.9 ± 0.5 versus 7.3 ± 0.6 ml/kg body weight, p=0.0002). Patchy and peribronchially accentuated lung fibrosis developed, with lympho-plasmacellular infiltrates, thickened septa, and distortion of the regular alveolar structure (Fig. 3D). As a result, an increase in tissue content, and a loss of airspace, were observed in the pepstatin A–treated animals (Fig. 3E). In parallel, a corresponding induction of ER stress, as indicated by alternative splicing of *Xbp-1* and Chop induction (Fig. S. 2C, D), was observed in pepstatin A–treated mice, alongside with a marked increase in cleaved Caspase 3 (Fig. S. 2D). Immunofluorescence localization of cleaved caspase and Chop demonstrated that in pepstatin A–treated mice the apoptotic signal was largely restricted to AEC2s (Fig. 3F).

Taken together these data demonstrate that *in vivo* inhibition of SP-B processing is sufficient to induce ER-stress and fibrosis in the mouse lung.

### Reactivation of Notch Signaling in IPF

To better understand which signaling pathways may underlie the regeneration of the alveolar epithelium in IPF, we performed transcriptome analysis on laser captured microdissected septa from donor (n=7) and IPF (n=10) patients. From IPF lungs, we collected normal appearing and fibrotic septa and compared each of them to septa from donor lung tissues. Thus, we generated two sets of pathways: those that were differentially regulated in normal appearing IPF regions versus donor septa, and those in fibrotic IPF regions versus donor septa. Notch signaling was differentially regulated in normally appearing septa of the fibrotic lung (Fig. 4A and Table S3) which were still populated by morphologically integer AEC2s. Because of this evidence of early abnormalities in Notch signaling in IPF and the prominent role of Notch in lung development and epithelial proliferation and differentiation, we subsequently focused our attention on Notch signaling.

**Fig. 4.**
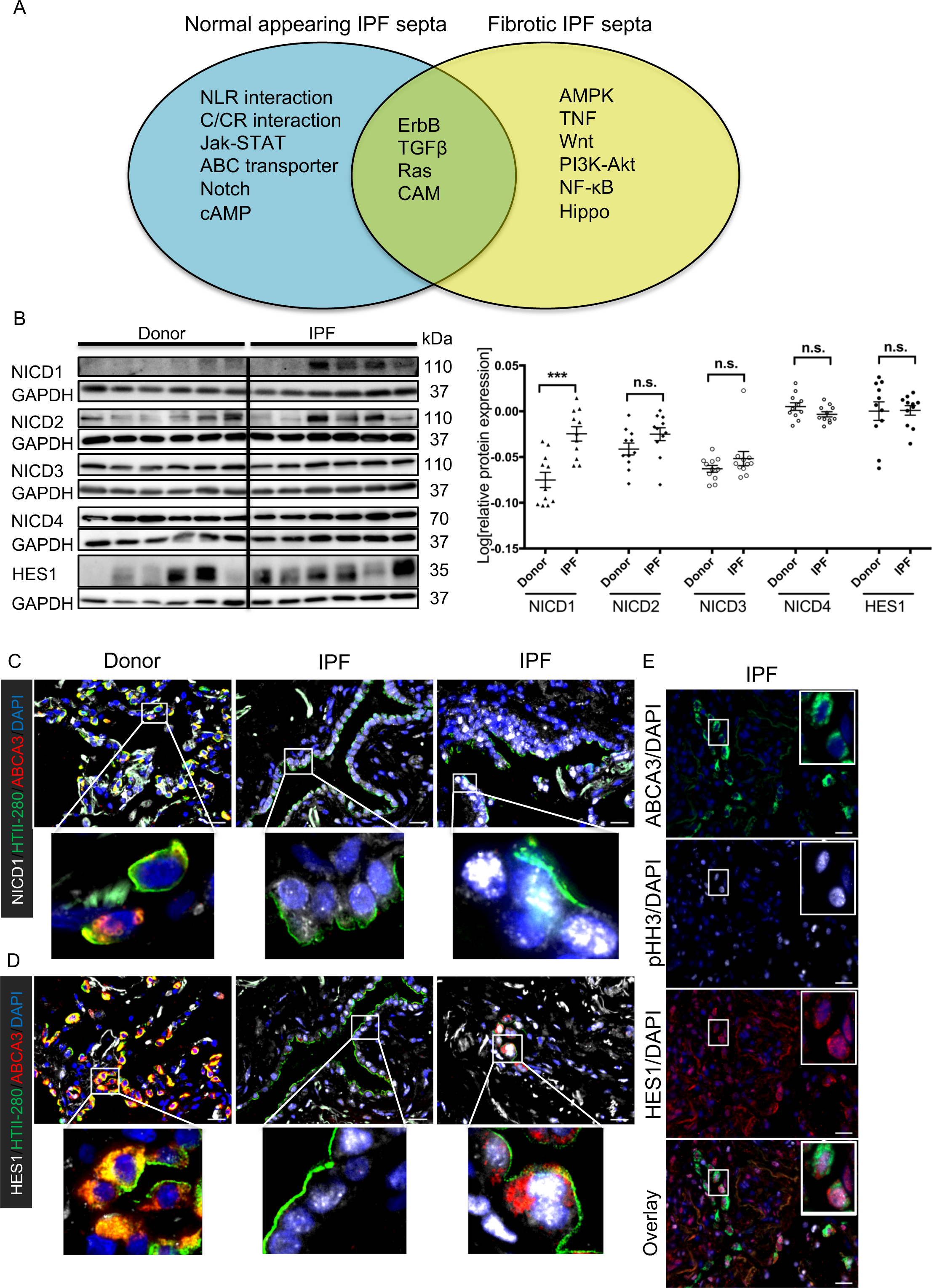
Notch1 signaling is upregulated in AEC2s from IPF patients. (A) Venn diagram displaying signaling pathways differentially regulated in normal appearing (blue) and fibrotic septa (yellow) from IPF patients (n=10) as compared with donor septa (n=7), with commonly regulated pathways in the intersection. (B) Western blot analysis and densitometric quantification of Notch receptor ICDs (NOTCH1–4) and HES1 expression in donor (n = 11) and IPF (n = 11) lung homogenates. GAPDH served as loading control. Data are presented as the mean ± SEM of log transformed densitometric values. ***p < 0.001, n.s. = not significant by unpaired Student’s t-test. (C), (D) Immunofluorescence localization of HTII-280 (green), ABCA3 (red), NICD1 (white in C), HES1 (white in D), in representative sections of donor and IPF (n=10 each) tissue samples. DAPI represents the nuclear counterstain. Scale bars, 20 μm. (E) Immunofluorescence localization of ABCA3 (green), HES1 (red) and phospho-HISTONE H3 (pHH3; in white) in representative sections of donor and IPF (n=10 each) tissue samples. DAPI represents the nuclear counterstain. Scale bars, 20 μm.

### NOTCH1 Activation in the Lungs of IPF Patients

To investigate Notch signaling activation in more detail, we analyzed Notch intracellular domains and Notch ligands at mRNA and protein levels in human IPF and donor lung homogenates. We found significantly increased levels of NICD1 receptor (Fig. 4B, n=11 donors and n=11 IPF patients) and DLL1 ligand (Fig. S. 3C n=6 donors, n=9 IPF) at the protein, but not the mRNA, level (Fig. S. 3A, B, n=5 donors and n=10 IPF) in peripheral lung tissues from patients with IPF. Otherwise, none of the other Notch receptors and ligands were differentially regulated in IPF lungs (Fig. 4, Fig. S.3C). Hes1, a key downstream target of all Notch receptors, was not significantly different between IPF and donor lung homogenates (Fig. 4B). However, immunofluorescence analysis of NICD1 and Hes1, a downstream target gene, revealed prominent nuclear immunoreactivity in AEC2s, identified by ATP-binding cassette sub-family A member 3 (ABCA3) or HTII-280 expression in both normal and fibrotic septa (Fig. 4C and D n=10 donors, n=10 IPF) and, to a certain extent, in other cell types (e.g., myofibroblasts and airway epithelial cells, endothelial cells; not shown in detail). In donor lungs, NICD1 was found in the nuclei of the bronchial epithelium and rarely and at much lower levels in AEC2s (Fig. 4C and data not shown). DLL1 staining was predominantly found in AEC2s of IPF lung (Fig. S. 3D). Finally, co-localization of the proliferation marker phospho-HistoneH3 with Hes1 indicated that AEC2s undergoing active Notch signaling are also proliferative (Fig. 4E n=10 donors, n=10 IPF). In addition, other, non-AEC2 cells showed nuclear expression of Hes1 and enhanced phospho-HistoneH3 staining. With regard to the alveolar expression of the other Notch receptors, we did not observe any differences between IPF and donors (Fig. S. 3E n=10 donors, n=10 IPF).

### Notch1 Signaling Activation in Bleomycin-Induced Fibrosis

To study Notch signaling in experimental lung fibrosis, we turned to the bleomycin-induced mouse model of lung fibrosis. On mRNA level, we did not observe meaningful changes in Notch receptors and ligands (Fig. 5 A, B n=4 mice per group). Upregulation of NICD1 and Hes1 proteins was encountered in lung homogenates during the proliferative (day 14) and fibrotic (day 21) phases of the response to bleomycin (Fig. 5C n=4 mice per group). At cellular level, AEC2s near fibrotic regions expressed high amounts of Notch1 and Hes1 in bleomycin-injured lungs at day 14, in contrast to the exclusive expression in the bronchial epithelium of saline-treated lungs (Fig. 5D). Notch1 upregulation was also observed in the non-epithelial compartment, but not in the conducting airways epithelium, in bleomycin-treated animals (data not shown). Importantly, isolated AEC2s from bleomycin-treated (n=5) mice at day 14 had increased proliferative activity that was paralleled by increased levels of NICD1 relative to saline controls (Fig. 5 E, n=4). Notch inhibition by the γ-secretase inhibitor DAPT decreased their proliferation (Fig. 5 F, G, n=3 mice per group). To further confirm the role of Notch signaling in lung fibrosis, we analyzed Notch1 and Hes1 expression in the above mentioned pepstatin A model. There, we also observed enhanced staining of AEC2 for Notch1 and nuclearization of HES1 in AEC2 (Fig. S. 2 E, n=4 mice per group), suggesting that Notch activation is a general feature of lung fibrosis.

**Fig. 5.**
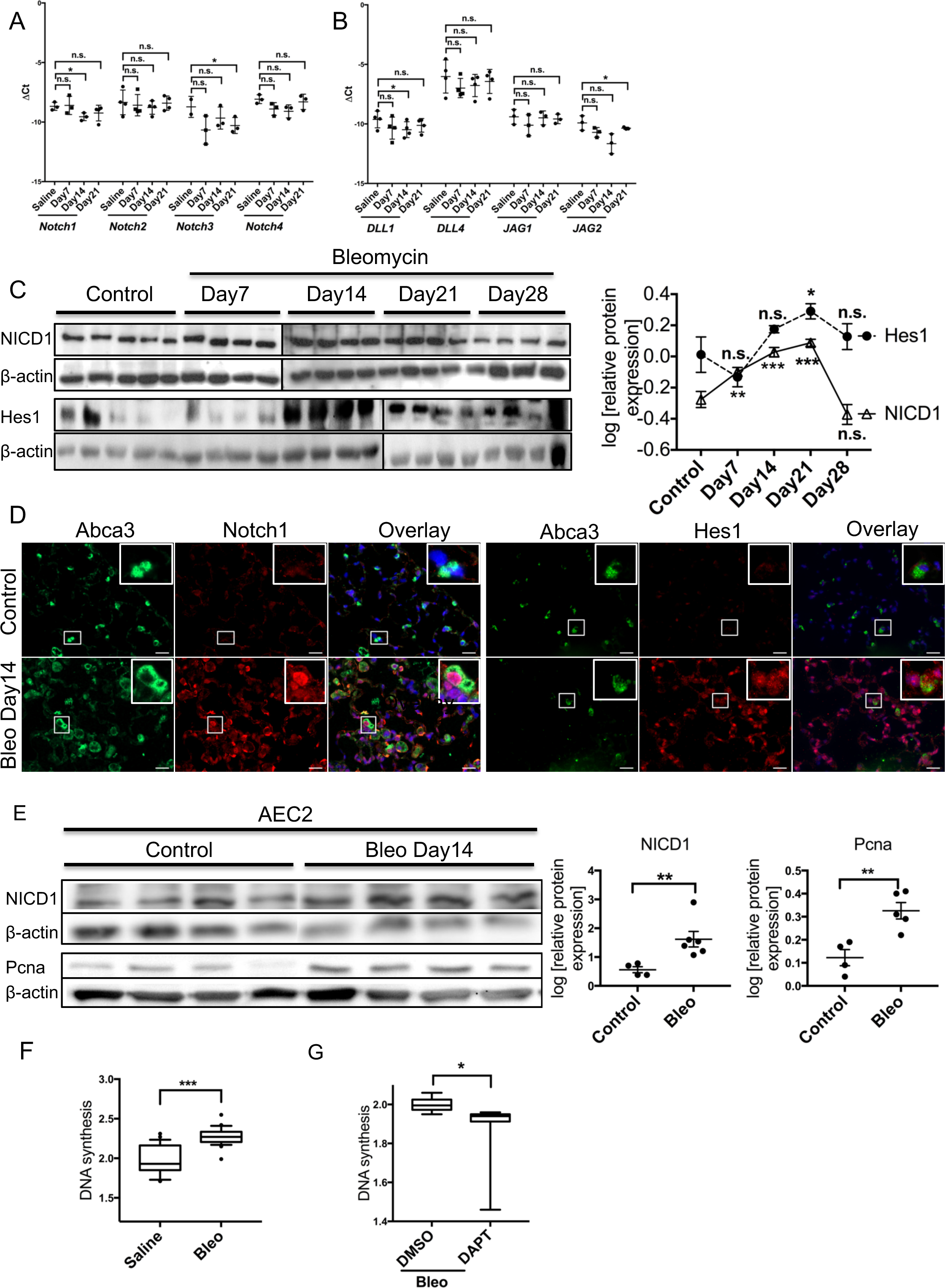
Notch1 signaling is upregulated in bleomycin-induced lung fibrosis. (A), (B) Expression of Notch receptor 1–4 (A) and ligand (Dll1, 4 and Jag1, 2; B) mRNAs in lung homogenates from bleomycin-treated C57BL/6N mice (day 7, 14, and 21) as compared with saline-treated controls(n = 4 animals per each group). Mean ΔCt (Cthousekeeping – Ctgene of interest) values ± SD are given. *p < 0.05, n.s. = not significant by unpaired Student’s t-test. (C) Western blot analysis and densitometric quantification of NICD1 and Hes1 in saline and bleomycin-treated C57BL/6N mice at day 7, 14, 21, and 28 (n=4 per group and time point). β-Actin was used as the loading control. Data are presented as the mean ± SEM of log transformed densitometric values. *p<0.05, **p<0.01,***p < 0.001, n.s. = not significant by unpaired Student’s t-test. (D) Immunofluorescence analysis of Abca3 (in green) and Notch1 or Hes1 (in red) in day14 saline- (Control, upper panel) or bleomycin-treated (Bleo Day14, lower panel) mice (n = 4). DAPI was used as nuclear counterstain. Scale bars, 20 μm. (E) AEC2s were isolated from day 14 saline- (n=4) or bleomycin- (n=5) treated mice and subjected to Western blot analysis (left) and densitometric quantification (right) of NICD1 and Pcna. β-Actin served as the loading control. Data are presented as the mean ± SEM of log transformed densitometric values. **p < 0.01 by unpaired Student’s t-test. (F) Cell proliferation was quantified in AEC2s isolated from day14 saline- or bleomycin-treated mice (n=4 each) by radioactive labeling of newly synthesized DNA with [H^3^]thymidine. Data are presented as the mean ± SEM. ***p < 0.001, by unpaired Student’s t-test. (G) AEC2s isolated from day 14 saline- or bleomycin-treated mice (n=3 each) were cultured for 24 h in the presence of DAPT (10 μM) or DMSO control. Proliferation was assessed by [H^3^]thymidine incorporation. Data are presented as the mean ± SEM. ***p < 0.001, by unpaired Student’s t-test.

### Impact of Notch Signaling on Alveolar Epithelial Cell Proliferation In Vitro

To further study the functional consequences of Notch1 upregulation in AEC2s, we overexpressed NICD1 in MLE12 cells and assessed the genome-wide transcriptome 12h (n=6 EV and n=6 NICD1), 24h (n=6 EV and n=6 NICD1), and 48h (n=7 EV and n=7 NICD1) after transient transfection. NICD1 overexpression resulted in a time-dependent increase in the number of genes involved in cell cycle regulation (Fig. S. 4A). Likewise, 5 of the 10 pathways identified here (Table S4) were also observed as being differentially regulated in normal or fibrotic laser capture microdissected septa from IPF lungs (Fig. 4A - CAMs, Jak-STAT, C/CR interaction, ErbB, TGF-β), suggesting that Notch signaling is upstream of these pathways. Functionally, overexpression of NICD1 in MLE12 cells resulted in significantly increased proliferation (Fig. S. 4B). In contrast, inhibition of Notch signaling by either siRNA-mediated down-regulation of *O*-fucosyltransferase (POFUT1) or DAPT treatment, resulted in a significant suppression of proliferation in both MLE12 cells (Fig. S. 4 C, D) and primary AEC2s obtained from mouse lungs 14 days after bleomycin treatment (Fig. 5G). Thus, modulation of Notch1 signaling regulates AEC2 proliferation via transcriptional regulation of cell cycle genes in the lung.

### AEC2-Specific NICD1 Overexpression In Vivo Leads to Lung Fibrosis

To study in vivo the consequences of epithelial upregulation of Notch1, we used doxycycline (Dox)-inducible, AEC2 specific NICD1-IRES-nEGFP overexpressing transgenic mice, to which we further refer to as NICD1 mice ^27 28 29^. Transgene induction was variable, leading to various levels of Cre and NICD1 (nEGFP) induction after 2 (n=8 mice) and 4 (n=14 mice) weeks of Dox treatment (Fig. S. 5 A). For this study, only mice with robust expression of both Cre and NICD1 (nEGFP) transgene (identified by an asterisk in Fig. S. 5 A n=3 no Dox, n=3 Dox 2 weeks, and n=4 Dox 4 weeks) were selected for further analysis. Transgene activation was also confirmed by an increase in Hes1 expression in AEC2s in a Dox-dependent manner (Fig. S. 5 B).

Histological assessment of the NICD1 transgenic mice sacrificed after 2 and 4 weeks of Dox treatment revealed an obvious fibrotic phenotype: patchy subpleural or hilar histological appearance of septal thickening, intra-alveolar infiltrates and increased matrix deposition (Fig. 6 A), reminiscent of some of the histopathological criteria of human UIP (usual interstitial pneumonia). Accordingly, in fibrotic areas of Dox-treated NICD1 transgenic mice, the septal thickness and the alveolar mean linear intercept were increased as compared with untreated controls (Fig. 6 B). Further immunofluorescence staining showed that in the fibrotic areas, but not in the non-fibrotic areas, of the Dox-treated transgenic mice, there was increased collagen and vimentin expression, along with enhanced Ki67 staining, reflecting increased proliferation in these areas (Fig. 6 C). Likewise, in lung homogenates, extracellular matrix and proliferation markers were significantly increased after 2 (collagen 1, αSMA) and 4 (vimentin, PCNA) weeks of Dox treatment (Fig. 6 D). Thus, distal lung epithelial overexpression of NICD1 results not only in proliferation of AEC2s, but also in mesenchymal activation and development of lung fibrosis.

**Fig. 6.**
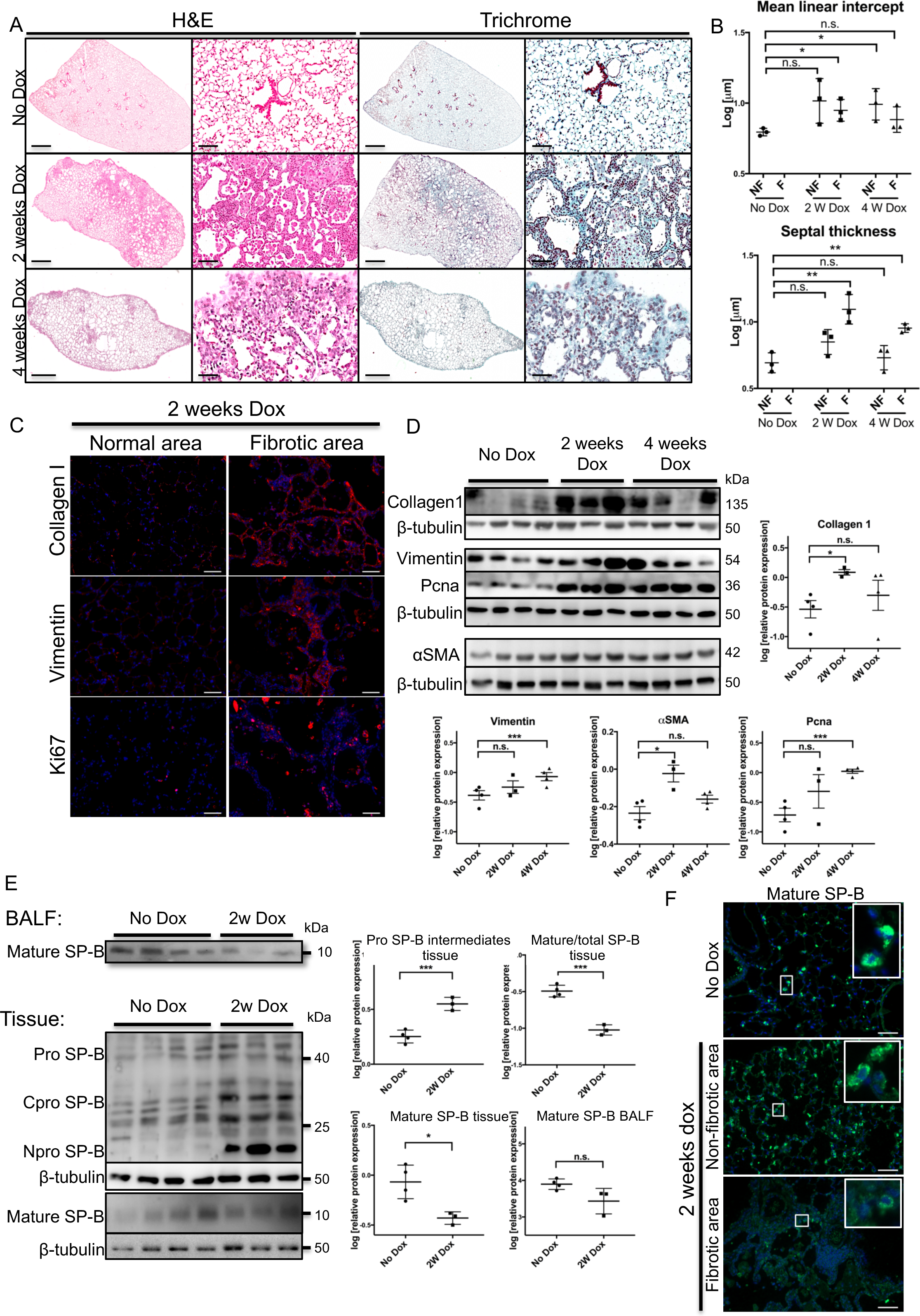
AEC2-specific NICD1 overexpression in vivo leads to lung fibrosis. (A) Representative images of H&E and Masson’s trichrome staining of NICD1 mice untreated (n=3) or doxycycline-fed for 2 (n=3) or 4 weeks (n=3). Scale bars in low-magnification images, 1 mm; in high-magnification images, 50 μm. (B) Morphometric analysis of septal thickness and alveolar mean linear intercept of mice as described in (A). Fibrotic areas (F) and non-fibrotic areas (NF) were analyzed separately. Data are presented as the mean ± SEM of log transformed densitometric values. *p < 0.05, **p < 0.01, n.s. = not significant by unpaired Student’s t-test. (C) Immunofluorescence analysis of Collagen I, Vimentin, and Ki67 expression in normal (i.e., non-fibrotic) areas and fibrotic areas in NICD1 mice exposed to doxycycline for 2 weeks. DAPI was used as nuclear counterstain. Scale bar, 50 μm. (D) Lung homogenates from mice as in (A) were subjected to Western blot analysis (left) and densitometric quantification (right) of Collagen 1, Vimentin, PCNA, and alpha-smooth muscle actin (αSMA). β-Tubulin served as the loading control. Data are presented as the mean ± SEM of log transformed densitometric values. *p < 0.05, ***p < 0.001, n.s. = not significant by unpaired Student’s t-test. (E) Western blot analysis and densitometric quantification of lung tissues and BALF from NICD1 transgenic mice unexposed (No Dox; n=3) or exposed to Dox for 2 weeks (n=3) using antibodies against pro- and mature SP-B. β-Tubulin served as the loading control. In case of BALF, equivalent volumes were loaded. Data are presented as the mean ± SEM of log transformed densitometric values. *p < 0.05, ***p < 0.001, n.s. = not significant by unpaired Student’s t-test. (F) Immunofluorescence staining of mature SP-B (green signal) was performed on lung tissue from mice as in (A). DAPI was used as a nuclear counterstain. Scale bar, 50 μm.

### NOTCH Activation Causes De-Differentiation of AEC2s with Altered Surfactant Protein Processing

In many other systems Notch signaling regulates cellular differentiation, which in AEC2s mainly entails the synthesis, processing, secretion and re-utilization of pulmonary surfactant. As surfactant processing by AEC2s was deeply affected in human IPF, and its inhibition is sufficient to induce lung fibrosis, we analyzed the status of the surfactant processing compartment in the NICD1 transgenic mice. We observed the very same pattern of surfactant abnormalities as found in patients with IPF: reduction of mature SP-B in the lung tissue and bronchoalveolar lavage fluid (BALF), in addition to a concomitant accumulation of proSP-B forms (Fig. 6 E). Correspondingly, the ratio of mature to total SP-B was significantly decreased in Dox-treated mice indicating disturbed intracellular processing of SP-B (Fig. 6 E). Immunofluorescence analysis of the lungs of the Dox-treated NICD1 transgenic mice for mature SP-B expression showed that there was almost no mature SP-B left in the fibrotic areas, whereas in the lungs of the same animals, mature SP-B was easily detectable in the non-fibrotic areas (Fig. 6 F).

### Inactivation of Notch Signaling Restores AEC2 Differentiation and Reverses Lung Fibrosis

To better understand if blocking Notch signaling would improve the differentiation status of AEC2s and surfactant processing, we followed an *ex vivo* and an *in vitro* approach, in which Lysotracker incorporation was used to measure the size of the surfactant processing compartment ^25^. First, MLE12 cells were treated with DAPT or vehicle alone, and the intensity of Lysotracker uptake was evaluated by flow cytometry after 3 days of treatment. We observed a dose-dependent and significant increase in Lysotracker fluorescence intensity alongside reduced Hes1 and NICD1 protein levels in response to DAPT treatment (Fig. S. 6 A-C). Second, we used precision cut lung slices (PCLSs n=1 per condition) obtained from the lungs of IPF patients at the time of transplantation to determine the concentration of DAPT required for effective downregulation of Notch signaling (50µM; Fig. S. 6 D). In DMSO treated PCLSs, the Lysotracker signal progressively decreased over a period of 4 days, while DAPT treatment resulted in a great increase in Lysotracker uptake (Fig. 7 A). Lysotracker uptake was also quantified by flow cytometry (n=7 IPF patients) in the EPCAM^pos^ CD45^neg^ CD31^neg^ live population (the full gating strategy is shown in Fig. S. 6 F). Again, there was a non-significant increase in the number (Fig. S. 6 E) and a significant increase in the mean fluorescence intensity of Lysotracker^pos^ cells in response to DAPT treatment (Fig. 7 B), despite the fact that the overall epithelial cell composition after 4 days was comparable between DAPT- and DMSO-treated PCLSs (Fig. S. 6 E). In parallel experiments, treatment of IPF PCLSs with DAPT resulted in an increase in mature SP-B, but not SP-C staining (Fig. 7 C and Fig. S. 6 G), albeit the overall level of mature SP was lower than in the explanted lung at day 0 (data not shown). Finally, DAPT treatment of IPF PCLS (from n=5 IPF patients) led to a remarkable loss of trichrome staining (Fig. 7 D) and collagen 1 (Fig. 7 E). Thus, blockade of Notch1 signaling in IPF PCLS appears to improve the differentiated AEC2 characteristics with regard to surfactant synthesis and processing and reversed the fibrotic remodeling.

**Fig. 7.**
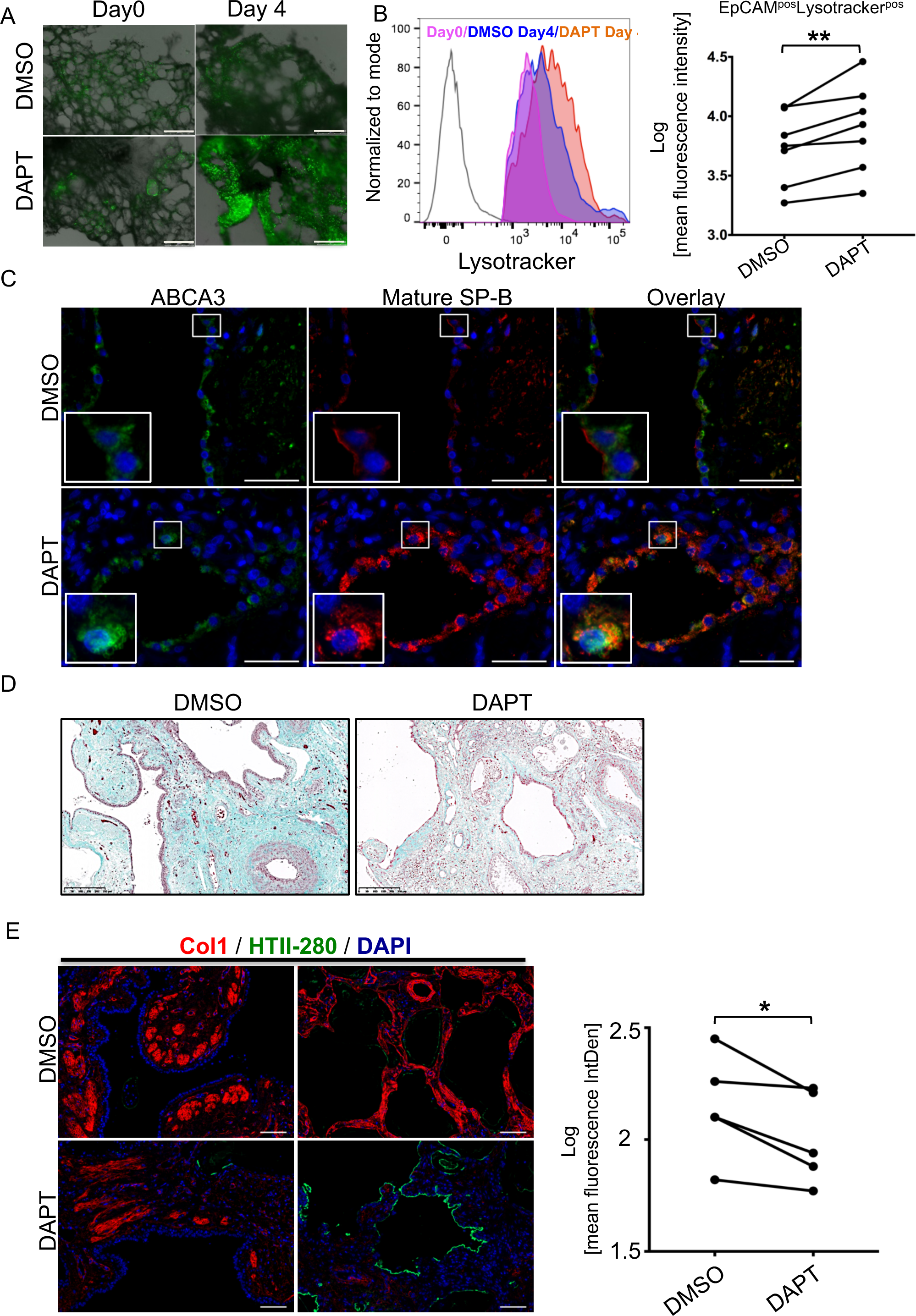
Inactivation of Notch signaling restores AEC2 differentiation and reverses lung fibrosis. (A) Live fluorescence imaging of human IPF PCLSs cultured in DMSO or DAPT (50 μM) for 4 days in the presence of Lysotracker Green. Representative overlaid phase contrast and Lysotracker Green images (n = 4 IPF patients) at Day 0 and 4 days (96 h). Scale bars, 500μm. (B) Human IPF PCLSs (n = 1-3 PCLS per condition from n=7 IPF patients) were cultured as in (A) and dissociated into single cells. Representative flow cytometry histograms of Lysotracker incorporation in the DAPI^neg^, CD45^neg^, CD31^neg^, EpCAM^pos^ population cultured for 4 days in the presence of DMSO (blue histogram) or DAPT (orange). Cells isolated at day0 from the lung of the same patient served as the control (pink). Unstained cells served as the negative control (unshaded histogram). Quantification of the mean fluorescence intensity of DAPI^neg^, CD45^neg^, CD31^neg^, EpCAM^pos^ Lysotracker Green^pos^ cells from n=7 patients is shown in the adjacent graph. Data in B (right panel) are presented as the mean ± SEM of either percentages of parent or log transformed mean fluorescent intensity. **p < 0.01 by paired Student’s t-test. (C) - (E) Human IPF PCLSs cultured for 4 days (3 PCLS per condition, n=5 patients) in the presence of DMSO or DAPT (50 μM) and stained (C) with an antibody specific for mature SP-B (red), shown is a representative image. AEC2s were identified by ABCA3 immunoreactivity (green signal). Scale bars, 20 μm; (D) PCLS stained with Masson’s trichrome staining. Scale bars, 200μm; (E) PCLS analyzed by immunofluorescence staining for Collagen 1(red), HTII-280 (green). DAPI was used for nuclear counterstain. Scale bars 50μm. Collagen1 staining intensity was quantified in the adjacent graph. Data are presented as the mean ± SEM of log transformed mean fluorescent intensity. *p < 0.05 by paired Student’s t-test.

## Discussion

Alterations in the pulmonary surfactant system have been previously described in the lungs of IPF patients ^24 30^ and in experimental models of lung fibrosis ^31 32^, but the underlying reasons and the clinical consequences have not been studied in depth. Here we show that human IPF lung is characterized by a pronounced increase in alveolar surface tension and alveolar collapse during expiration, which appears to be largely caused by a massive processing defect of the hydrophobic surfactant proteins SP-B and SP-C. Such increase in alveolar surface tension, which can be almost completely rescued *ex vivo* by addition of mature SP-B alone, must be considered to contribute to the two key pathophysiological changes in lung fibrosis: loss of pulmonary compliance (as suggested in animal models of lung fibrosis ^33,34,35,36^) and impaired gas exchange (as evident from V/Q assessment in human IPF ^20^ and experimental models ^35,36^ of lung fibrosis). It therefore remains an intriguing thought that therapeutic correction of the alveolar surface tension in IPF could result in improved lung mechanics or gas exchange or both, similarly to other clinical conditions with profound surfactant abnormalities such as the Infant ^37^ or the Adult ^38^ Respiratory Distress Syndrome. Moreover, in our study, experimental inhibition of surfactant processing, with far-reaching loss of mature SP-B in the mouse lung, was sufficient to cause overt lung fibrosis. Such observation fits well to the very rare cases of inherited SP-B deficiency with partial defects in SP-B production, in which the concerned children survive beyond the neonatal period and develop Interstitial Lung Disease (reviewed in ^39^). Although additional studies will be needed to explore this in more depth, one plausible explanation could be enhanced AEC2 injury, either through intracellular accumulation of unprocessed surfactant proteins and induction of ER-stress - as shown in here *in vitro* and *in vivo* - or enhanced stretch of AEC2 due to repetitive alveolar collapse ^40^, with secondary, paracrine, activation of fibroblasts. Another reasonable explanation could be stretch-induced, αvβ5 integrin – mediated, direct activation of TGF-β in the neighbouring mesenchyme, as suggested previously ^41^. As a result of less retractive forces of neighbouring tissue in the subpleural, and a higher transpulmonary pressure gradient in an otherwise incompliant lung in the subpleural and basal areas, it appears reasonable that a permanent and uniform elevation of alveolar surface tension in the lung may promote evolution of fibrotic changes preferential in the subpleural and basal parts of the lung, a pertinent finding in IPF ^23^.

Importantly, we identified Notch signaling as major pathway differentially regulated in IPF, and demonstrate that Notch1, in particular, is activated in AEC2s of IPF patients and bleomycin-injured mouse lungs. In both cases, its expression correlated with increased proliferation, and blocking Notch signaling dampened proliferation of alveolar epithelial cells isolated from day 14 bleomycin treated mice and MLE12 cells. AEC2-specific overexpression of NICD1 in the mouse lung recapitulated major aspects of human IPF: an irregular, patchy and in part subpleural distribution, increased collagen deposition and comparable disturbances in the surfactant system. Most importantly, we show that blocking Notch signaling in human IPF PCLS has major therapeutic benefits, namely reduction in collagen deposition and improved differentiation of AEC2, including restoration of mature SP-B.

Notch signaling plays a major role in lung development ^42^. Its expression starts with lung bud formation and follows the proximo-distal axis, suggesting it may control cell fate decisions along this axis ^42–45^. In this regard Notch1 deeply affects the differentiation of epithelial-specific lineages in the airway epithelium. ^19,46,47,18,48^. However, its downregulation is necessary to allow specification of the alveolar fate ^43,18^. In a previous study ^49^ overexpression of NICD1 in adult mouse AEC2s using the CCSPrtTA driver line led to AEC2 proliferation in the first 14 days, followed by extensive compensatory apoptosis. In our work, we used the SPCrtTA driver line to overexpress a mouse NICD1 construct lacking the PEST domain, which is responsible for its degradation ^27^. Similarly, we see an increase in proliferation after 14 days of transgene induction, but at the same time we find major defects in SP processing and fibrosis. Lung fibrosis was not described in the study by Allen et al. ^49^ and this may be due to the different level and spatial localization of the NICD1 inherent to the transgenic mouse models used. We therefore think that there is a threshold level and spatial distribution of Notch activation beyond which lung fibrosis develops.

In conclusion, we propose a model in which Notch1 regulates the balance between the proliferative and differentiated functions of AEC2s in the human lung (see Fig. 8). Limited injury to the alveolar compartment results in Notch1 activation in AEC2s, which undergo a transient de-differentiation, with incomplete surfactant processing, in order to allow the activation of their progenitor function and regenerate the damaged epithelium. Repetitive alveolar injury as in IPF, however, results in massive proliferation and de-differentiation of AEC2s by Notch1 activation. The resulting loss in mature SP has severe consequences on alveolar homeostasis, leading to a more general increase in surface tension, alveolar collapse and establishes a vicious cycle of injury and repair leading to lung fibrosis. Hence, regional and timely limited Notch activation, with a temporary atelectasis of one lung region (as in case of lobar pneumonia), may allow complete regeneration of the alveolar epithelium in absence of biomechanical forces. In contrast, persistent and global Notch activation, as in IPF, may be detrimental and may contribute to fibrosis development. Thus, we propose Notch signaling is a promising therapeutic target to re-establish alveolar epithelial homeostasis and interfere with initiation and progression of fibrosis.

**Fig. 8.**
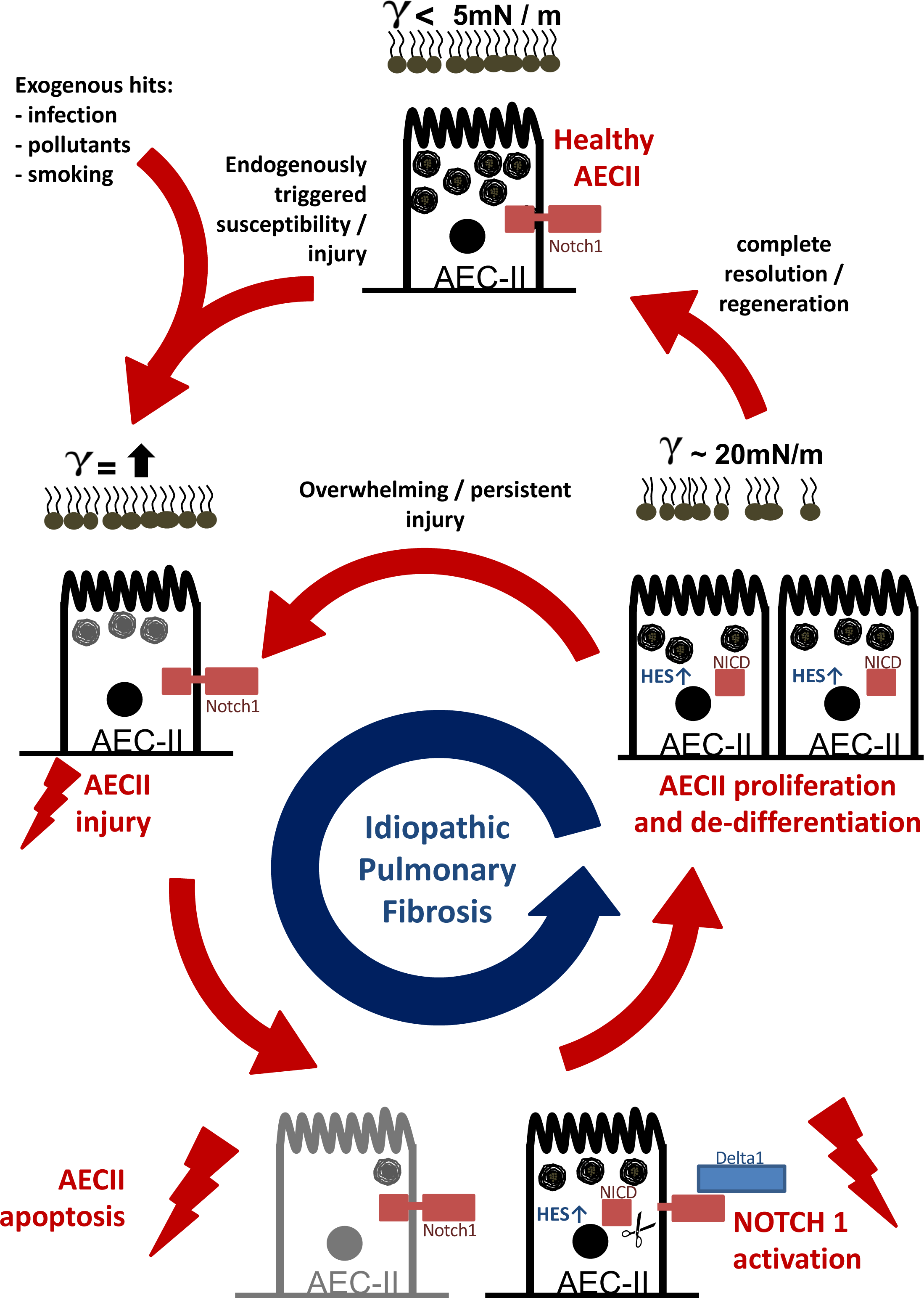
Summary of the data. Vicious cycle of alveolar epithelial injury, Notch activation, epithelial proliferation and de-differentiation, increased alveolar surface tension and collapse, and alveolar stretch-induced additional cell injury in Idiopathic Pulmonary Fibrosis. For details see Discussion.

## Material and Methods

### Study design

The objective of the study was to identify the role of Notch1 driven AEC2 de-differentiation and loss of surfactant processing in the pathogenesis of IPF. To that end we employed patient-derived materials (explanted lung, isolated cells, BALF, blood), transgenic and non-transgenic mice, primary and transformed cell lines as outlined below.

We included IPF or donor materials in our experiments. All IPF diagnoses were made according to the American Thoracic Society (ATS) / European Respiratory Society (ERS) consensus criteria (Raghu et al, 2011), and a usual interstitial pneumonia (UIP) pattern was shown to be present in all IPFLTX and IPFVATS patients. N numbers were dependent on sample availability (in case of PCLS experiments requiring fresh, living tissue) and on previous experiences made in human IPF lung tissues samples and cells ^50–52^ and are specified for each experiment.

We used bleomycin treatment of C57/B6N mice as a model of lung fibrosis. For these experiments we included mice between 12-16 weeks of age, weighting 18–22 g. In order to inhibit surfactant protein processing, Pepstatin A or solvent alone were administered intratracheally as aerosol to mice of similar age and weight. To study the effects of NICD1 overexpression in AEC2s, SPC-rtTA × tetO-Cre × NICD fl/fl triple-transgenic mice were fed doxycycline-enriched or regular standard lab chow. Animals weight and activity were monitored daily throughout the experiment. At each time point, mice were euthanized, lung compliance was measured, and tissue was harvested for protein, mRNA isolation and immunofluorescence analysis. N numbers in these experiments were determined based on our previous experience with the model and the readouts ^53,54^. In the case of the NICD1 we excluded mice were appropriate levels of transgene induction was not achieved (see Fig. S. 5C).

### Patient and Control Groups

Phenotyping and acquisition of biomaterials from patients were undertaken as a part of the European IPF Registry and Biobank (www.pulmonary-fibrosis.net) and the University of Giessen Marburg Lung Center Biobank. Both study protocols were approved by the Ethics Committee of the Justus-Liebig-University School of Medicine (No. 31/93, 29/01, and 111/08). Informed consent was obtained in written form from each subject (see overview of included subjects in Supplemental Table 1).

### High-Resolution Computed Tomography (HRCT) Analysis

HRCT scans were obtained during routine diagnostics. Overall, scans from 33 patients were collected (24 IPF; five COPD; four controls diagnosed with obstructive sleep apnea syndrome, pulmonary hypertension, congestive heart disease, and absence of a structural lung disease). HRCT scans were analyzed using Dicom Works 1.3.5b (2003, Philippe Puech, Loic Boussel) to obtain raw Hounsfield Unit (HU) distributions at the apical lung level, at the level of the tracheal bifurcation, and at the level of the basal lung either during inspiration or expiration. PROC MEAN was used to calculate mean lung attenuation from HU distributions using SAS 9.4 (Cary, USA). To demonstrate putative alveolar collapse, CT scans during inspiration and expiration were segmented using Dicom Works and were color-coded as described ^55^ using the GNU Image Manipulation Program: hyperinflated = – 1000 to −901 = red; normally aerated = −900 to −501 = blue; poorly aerated = −500 to −101 = yellow; non-aerated = HU −100 to 100 = green). In addition, individual sections were hand-selected, transferred and analysed for HU distribution. All data from all patients per group were pooled and are given as mean densitogram for the 3 patient categories and inspiration versus expiration and according to the percentage of voxels belonging to the above mentioned density categories (as mean +/-SEM). Significance of these categorial changes was assessed using the Wilcoxon-Mann-Whitney Test). Finally, the difference between the mean lung attenuation during inspiration and expiration was used to determine the density shift that occurs from inspiration to expiration. This data was provided as median +/-interquartile range and significance level was assessed by ANOVA.

*Immunofluorescence, immunohistochemistry, Western blotting, flow cytometry, cloning, genotyping and microarray analysis* were performed using standard protocols. Detailed procedures and antibodies used throughout the study are outlined in Supplementary Materials section and tables S2 and S5.

### Animal Studies

All animal studies were performed in accordance with the Helsinki convention for the use and care of animals and were approved by the local authorities at Regierungspräsidium Giessen GI20/10 Nr. 99/2010 (Notch mice), GI20/10 Nr. 109/2011 (Bleomycin), and GI20/10 Nr. 74/2007 (Pepstatin).

### Bleomycin Model of Lung Fibrosis and Chronic Pepstatin Administration

C57BL/6N mice (Charles River Laboratories, Sulzfeld, Germany) between 12-16 weeks of age, weighting 18–22 g, were used in all experiments. Bleomycin (Hexal) in 0.9% sodium chloride (saline) at a dose of 5 U/kg body weight was administered intratracheally as previously described ^53^. The specific napsin A inhibitor pepstatin A (Applichem, Darmstadt, Germany; 75 mg/kg body weight) was administered daily over 4 weeks as an intratracheal bolus. At indicated time points mice were sacrificed and their lungs harvested for further analysis.

### Isolation of mouse AEC2s and cell culture

AEC2 isolation was performed using dispase digestion followed by magnetic beads separation of CD16/32, CD45, CD31 negative cells as previously described ^56^. The AEC2s were plated in 48-well plates (150,000 cells/well) or used for cytospin preparations. Cells were grown up to 72 h in DMEM supplemented with 1% FBS, 1% penicillin/streptomycin, and 10 mM HEPES. Murine alveolar MLE12 cells (ATCC CRL 2110) were grown according to the ATCC suggested conditions. All cultures were maintained in a humidified atmosphere with 5% CO2 at 37°C.

### Isolation of human AEC2s

Human IPF or donor tissue were used for cell dissociation for flow cytometry. Thus, the tissue was weighed, chopped into small pieces of approximately 1-2 cm and further minced until a uniform granular suspension was obtained (approx. 1 mm pieces). Minced tissue was washed 3 times with RPMI and then resuspended in a solution of 10 % Dispase (Cornig) 30 μg/ml DNAse I (Sigma-Aldrich) in RPMI (Thermo Scientific) at a volume of 2ml Dispase solution per gram of tissue. Tissue was then incubated at 37°C on a rotating shaker for 2h. At the end of incubation time, the solution was pipetted up and down to maximize the cell release, strained through gauze, 100 μm, 70 μm and 40 μm cell strainer to obtain a single cell suspension. Cells were cryopreserved in 20% FBS (PAA) 5% DMSO (Sigma Aldrich) RPMI in the UGMLC Biobank until the time of the flow cytometry analysis.

### High-Precision Cut Lung Slices (PCLS) Generation and Culture

One segment of each explanted human IPF lung was filled with 1.5% Low Melt Agarose (Bio-Rad) at 37°C and allowed to cool on ice for 30 min. Blocks of tissue of were sectioned using a Vibrating Blade Microtome (Thermo Fisher) at a thickness of 500 μm. PCLSs were cultured for 4 days in RPMI (Gibco) supplemented with 10% human serum (Human Serum Off The Clot, Biochrom) and 1% penicillin/streptomycin in the presence of DAPT (10–50 μM in DMSO) or DMSO (1:1000; Sigma-Aldrich) with daily medium changes. For dissociation we used a similar protocol to that described for human AEC2 isolation.

### Data and software availability

All data are available upon request to the lead author. The microarray data obtained from microdissected human lung tissue and the microarray data from NICD1-transfected MLE12 cells have been deposited in the Gene Expression Omnibus (https://www.ncbi.nlm.nih.gov/geo/, accession Nos. GSE68239 and GSE95856, respectively).

### Statistics

For the statistical comparison of differences between two groups (IPF vs. Donor/HV, IPF vs. COPD, COPD vs. HV, pepstatin A–treated mice versus controls), the Student’s t-test was applied on log-transformed values. Comparison of more than two groups (HR-CT quantification) was performed using a one-way ANOVA followed by Tukey’s multiple comparisons of means. For the treated vs untreated PCLS (DAPT treatment), paired t-test was applied. For statistical analyses, the software GraphPad Prism version 5.02 was used. A p-value < 0.05 was considered statistically significant. Detailed explanation of microarray data analysis, quantitative lung morphometry and High-Resolution Computed Tomography (HRCT) analysis is found in Supplemental Materials.

## Supplementary Materials

### Supplementary Materials and Methods

#### Supplementary Figures

Fig. S1. Defective surfactant processing in IPF.

Fig. S2. Defective surfactant processing in pepstatin A-treated mice

Fig. S3. Notch signaling in human donor and IPF lungs

Fig. S4. Consequences of NICD1 upregulation in vitro

Fig. S5. Transgene expression in NICD1 overexpressing mice

Fig. S6. In vitro Notch inhibition restores differentiation of AEC2s

#### Supplementary Tables

Table S1. Basic Characteristics of the Patients Involved in our Studies

Table S2. *SFTPB, SFTPC*, and *NAPSA* Genotyping in IPF Patients and Healthy Controls

Table S3. Top Ten Signaling Pathways That Are Differentially Regulated in the Normal Appearing Septa from IPF Patients Versus Donor Septa and in Fibrotic Septa from IPF Patients Versus Donor Septa

Table S4. Differentially Regulated Cellular Processes in MLE12 Cells after 24 and 48 h of NICD1 Upregulation

Table S5. Antibodies used in this study

Table S6. Chemicals, peptides and recombinant proteins used in this study.

Table S7. Commercial assays used in this study

Table S8. Oligonucleotides used in this study

## Acknowledgments

The authors are grateful for the superb technical assistance of Silke Händel and Stefanie Hezel and for the valuable advice of Tilman Borggrefe (Department of Biochemistry, Faculty of Medicine, Justus-Liebig-University Giessen). Funding: This study was supported by grants of the German Ministry of Science and Education (“German Center for Lung Research”, DZL), the European Commission (“European IPF Network”, FP7), the German Research Council (“Excellency Cluster Cardio-Pulmonary System”; KFO 309).

## Author Contributions

The study was designed and supervised by AG, WS, TB, SB and OE. RW, MKorfei, CR, KP, PM, MKoenigshoff, EA, OK, and IS conducted experimental work. WK prepared and provided human lung materials. LF provided pathological evaluation of human and animal data. Animal experiments were primarily conducted by IH. Transcriptome analysis was undertaken by JW. HW conducted the sequencing work. Human CT analysis was performed by DvdB. AG and RW assembled the figures and wrote the manuscript.

