## Supplementary figures and images for "Restored alveolar epithelial differentiation and reversed human lung fibrosis upon Notch inhibition"

### Supplemental Figure 1

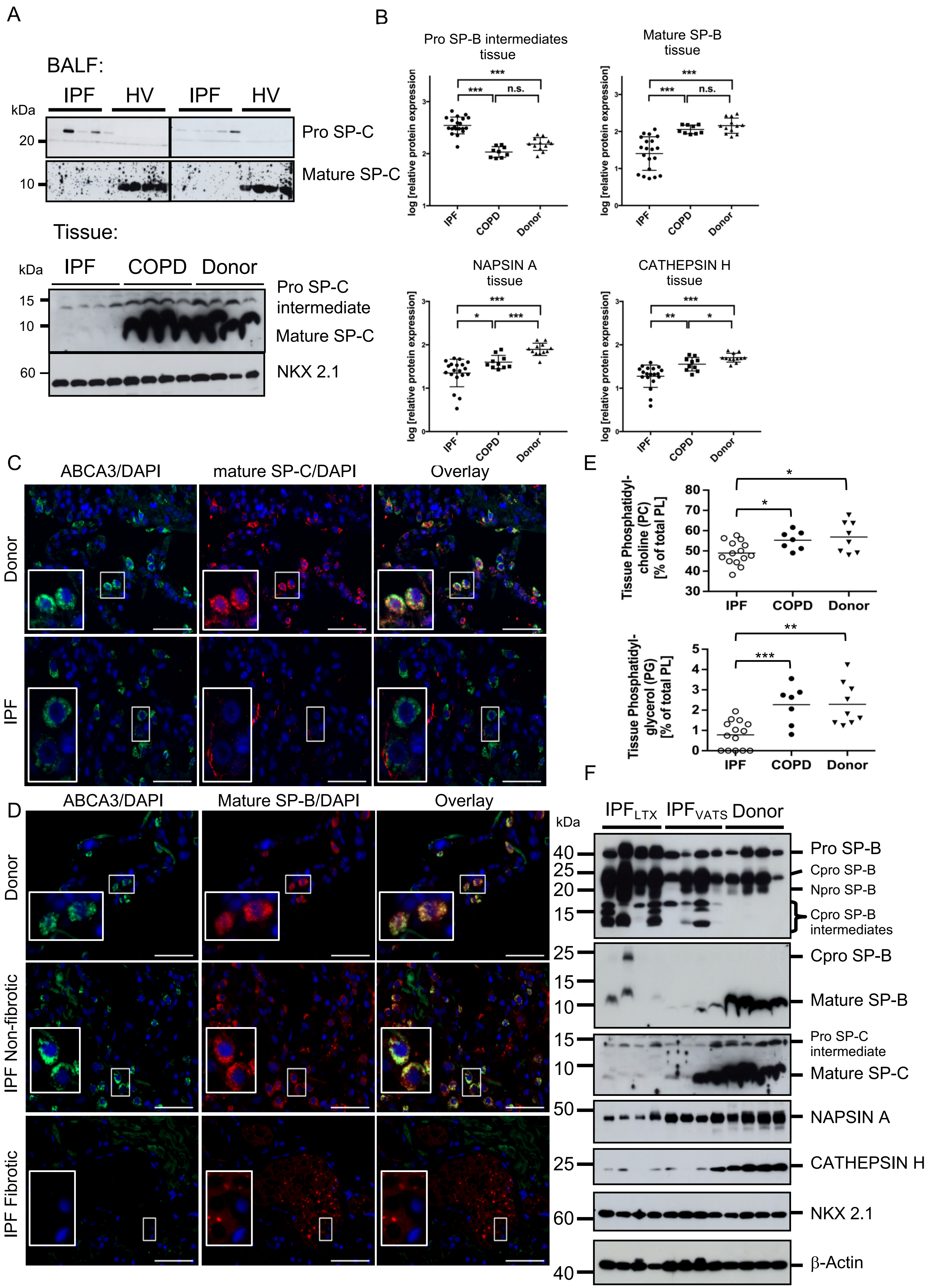

### Supplemental Figure 2

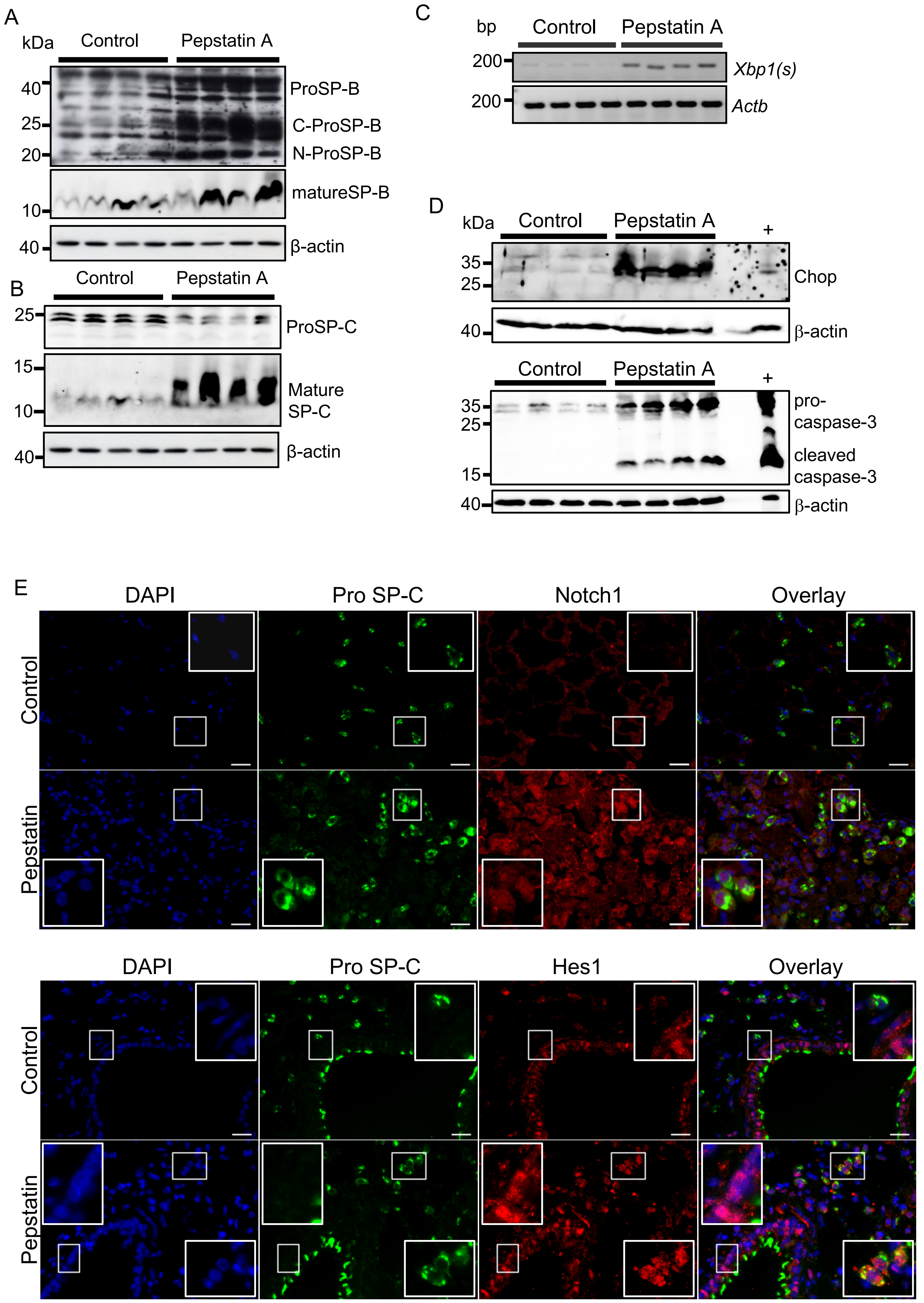

### Supplemental Figure 3

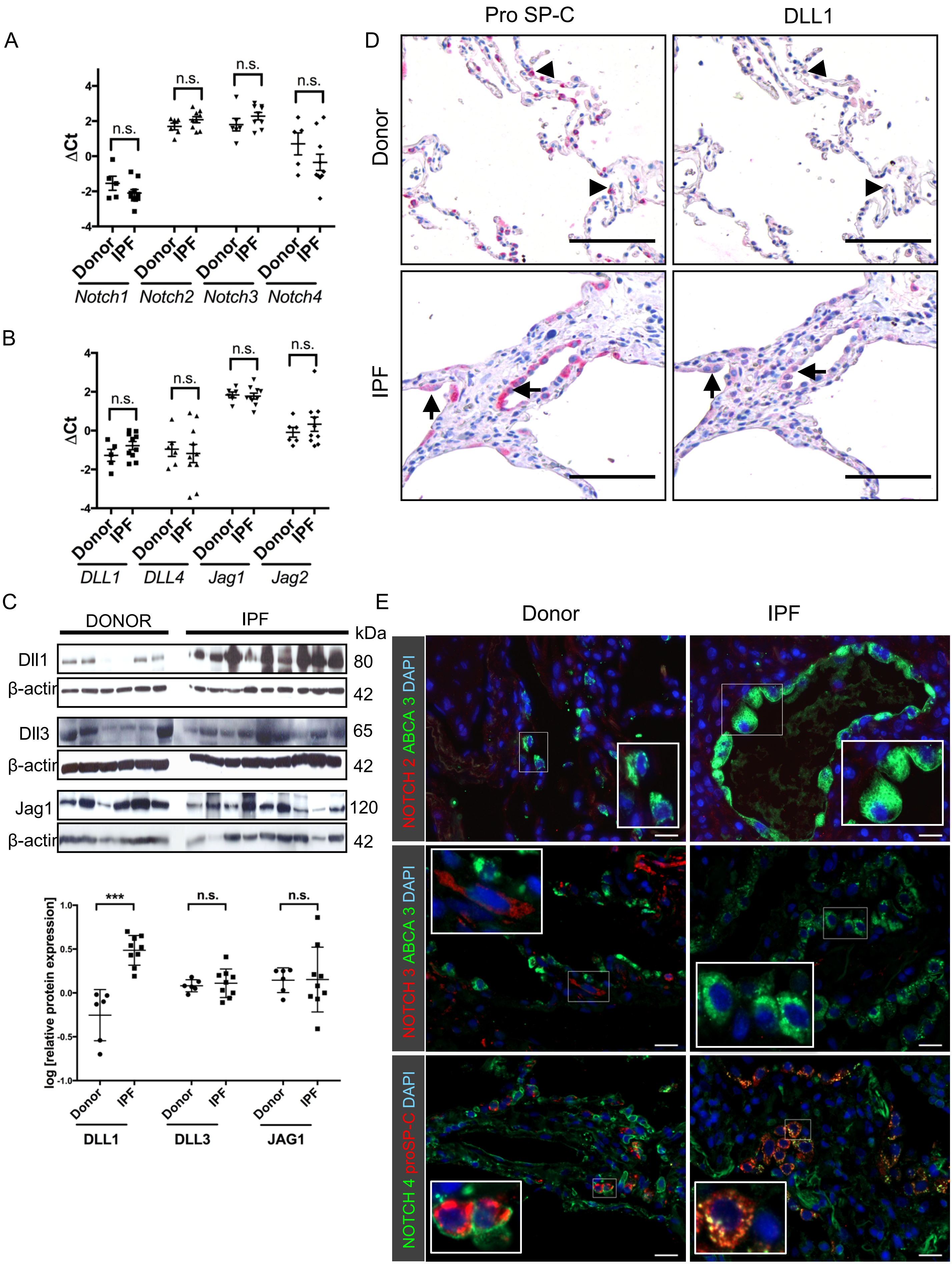

### Supplemental Figure 4

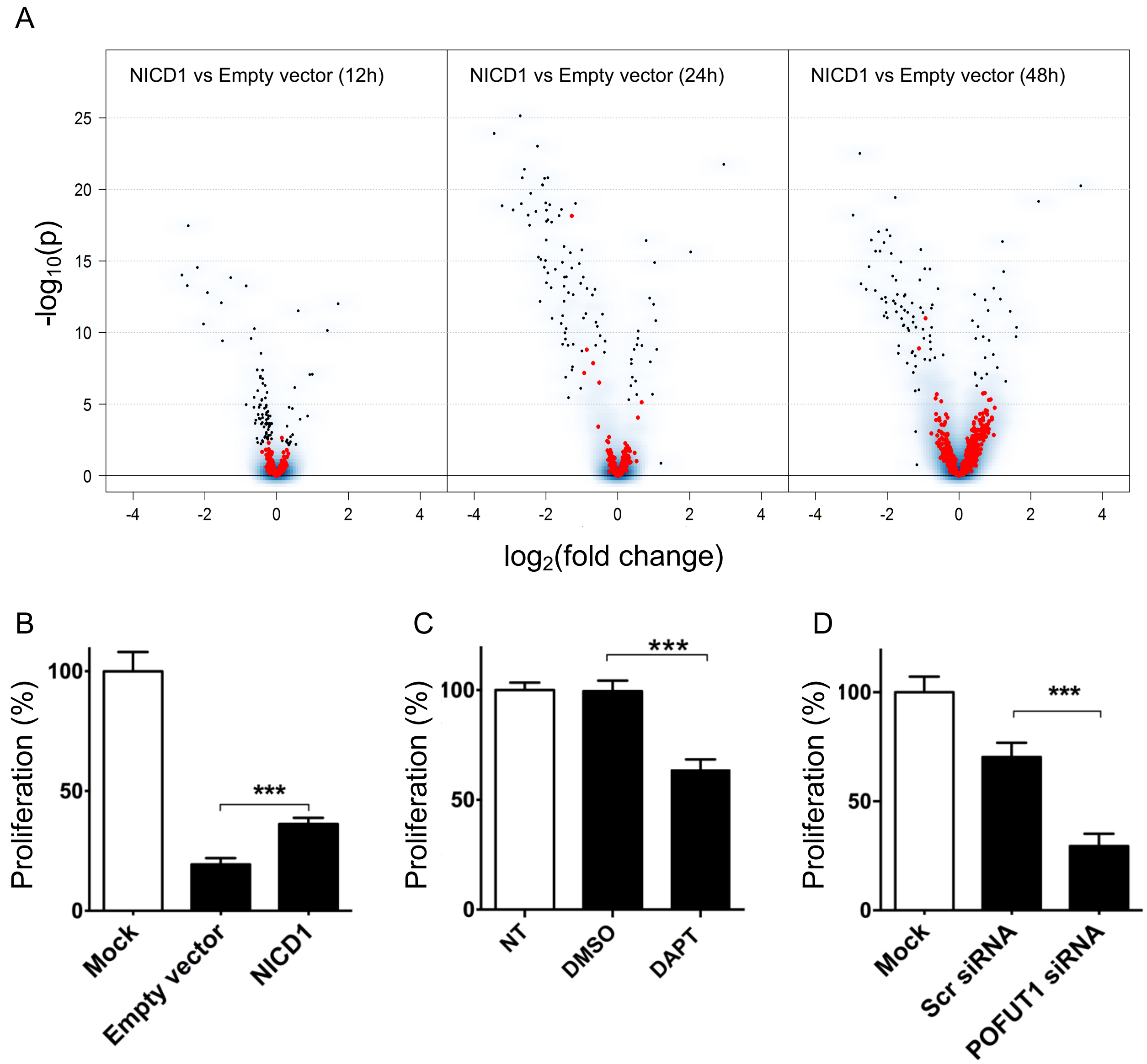

### Supplemental Figure 5

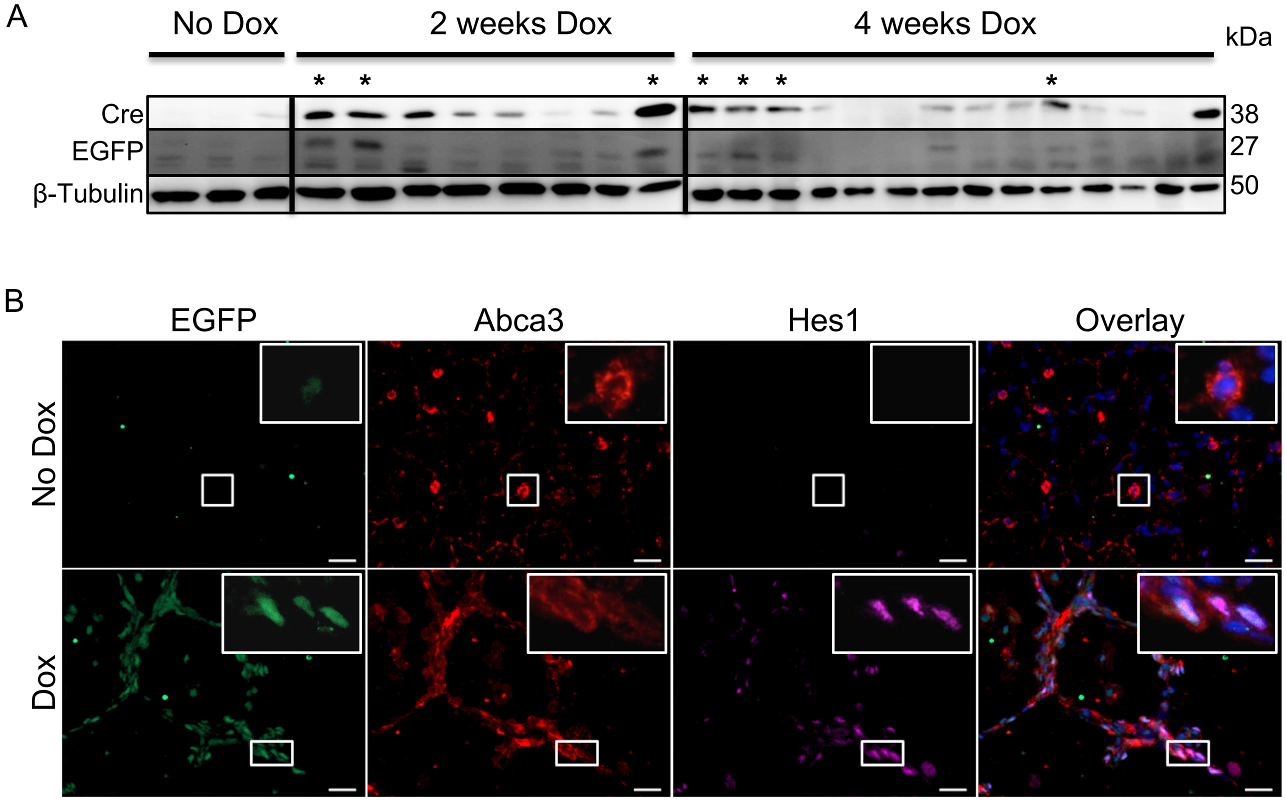

### Supplemental Figure 6

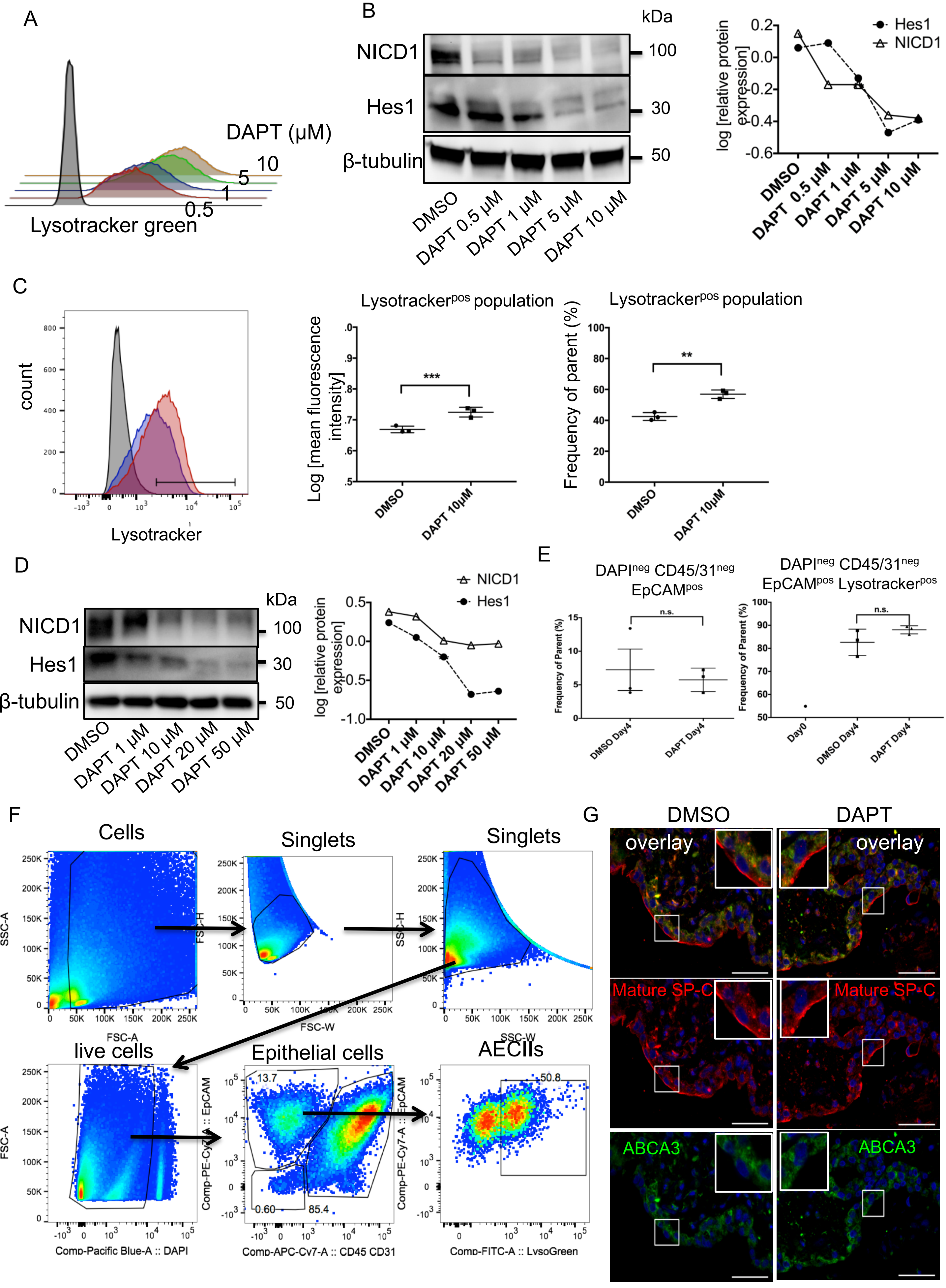
