## Supplemental Figure Legends for "Restored alveolar epithelial differentiation and reversed human lung fibrosis upon Notch inhibition"

**Fig. S1.** Defective surfactant processing in IPF. (A) Western blot analysis of BALF (upper panel) from IPF patients (n = 10) and healthy volunteers (n = 8) for 21-kDa proSP-C and mature SP-C and of peripheral lung tissue (lower panel) from IPF (n = 20), COPD (n = 9), and human donor (n = 12) lung tissues for proSP-C intermediates, mature SP-C and NKX.2. (B) Densitometric quantification of mature SP-B western blots as presented in Fig. 2A. Data are given as mean ± SEM of log transformed densitometric values. *p<0.05, **p<0.0, ***p < 0.001, n.s. = not significant by unpaired Student´s t-test. (C) Representative immunofluorescence staining for ABCA3 (green) and mature SP-C (red) in human lung tissue (n = 9 donors, n = 10 IPF patients). Scale bars, 20 $\mu$m. (D) Representative immunofluorescence staining for ABCA3 (green) and mature SP-B (red) in human lung tissue (n = 9 donors, n = 10 IPF patients). Scale bars, 20 $\mu$m. (E) Tissue stores of phosphatidylcholine (PC) and phosphatidylglycerol (PG) in IPF (n = 14), COPD (n = 7), and healthy donor (n = 9) lung tissues. The relative content of phosphatidylcholine and phosphatidylglycerol is given as a percentage of all phospholipids (PLs). Data are presented as mean values. *p < 0.05, **p < 0.01, ***p < 0.001, n.s. = not significant, by Student's t-test. (F)Western blots of lung tissue homogenates obtained from 4 IPF LTX explant tissues and 4 IPF open lung biopsies and 4 healthy donor tissues for proSP-B, mature SP-B and SP-C, NAPSIN A, CATHEPSIN H, and NKX2.1. β-ACTIN served as the loading control.

**Fig. S2.** Defective surfactant processing in pepstatin A-treated mice. (A) Representative immunoblot analysis for pro and mature SP-B (A) in lung tissue homogenates from vehicle (Control) or pepstatin A (Pepstatin A) treated mice as described in Figure 3 (n = 4 per group). β-actin was used as loading control. (B) Representative immunoblot analysis for pro and mature SP –C in lung tissue homogenates from the same mice (n=4 per group). β-actin was used as loading control. (C) RT-PCR analysis of spliced xbp-1 (*xbp1(s)*) in lung tissue homogenates from the same mice (n = 4 per group). *Actb* was used as house-keeping gene. (D) Representative immunoblot analysis of Chop (upper panel) and cleaved Caspase-3 (lower panel) in lung tissue homogenates from the same mice. As an additional control one lane (+) was added, which contained cell lysates from staurosporine treated murine, alveolar epithelial cell line (MLE12). (E) Representative immunofluorescence staining for proSP-C (green) and Notch1 in lung tissue sections from pepstatin A– and vehicle-treated mice. Scale bars, 20 $\mu$m. (F) Representative immunofluorescence staining for proSP-C (green) and Hes1 in lung tissue sections from pepstatin A– and vehicle-treated mice. Scale bars, 20 $\mu$m

**Fig. S3.** Notch signaling in human donor and IPF lungs. (A), (B) Quantitative real-time (q)PCR analysis of Notch signaling receptors *NOTCH1–4* and ligands (*DLL1,4* and *JAG1,2*) from donor (n = 5) and IPF (n = 10) lung homogenates. Data are represented as mean± SEM of $\Delta$Ct ($Ct_{house\ keeping}$ - $Ct_{gene\ of\ interest}$), n.s. = not significant by unpaired Student´s t-test. (C) Western blot analysis of DLL1, DLL3 , and Jagged1 in donor (n = 6) and IPF (n = 9) lung homogenates (upper panel). The lower panel shows quantification of the western blots. Data are presented as the mean ± SEM of log transformed densitometric values. ***p < 0.001, n.s. = not significant by unpaired Student´s t-test. (D) Immunohistochemistry localization of proSP-C and DLL1 in adjacent serial sections from donor and IPF patients. Arrowheads (donor) and arrows (IPF) identify AEC2s with immunoreactivity for proSP-C (left panel) and DLL1 (right panels, all in red). (E) Representative images of immunofluorescence analysis of 10 donor and 10 IPF lung tissues for ABCA3 (green), NOTCH2 (red) and DAPI (blue, upper row); ABCA3 (green), NOTCH3 (red) and DAPI (blue, second row); NOTCH4 (green), ABCA3 (red) and DAPI (blue, lowest row).

**Fig. S4.** Consequences of NICD1 upregulation in vitro. (A) Microarray data from MLE12 cells transiently transfected with empty vector or NICD1 and analyzed 12 (n=6 EV and n=6 NICD1), 24 (n=6 EV and n=6 NICD1), and 48 h (n=7 EV and n=7 NICD1) after transfection. Volcano plots show differentially regulated cell cycle genes (red dots) overlaid with all up and down-regulated genes (black dots) at each time-point. (B) Cell proliferation in NICD1-transfected, empty vector–transfected, and mock-transfected MLE12 cells was assessed by radioactive labeling of newly synthesized DNA with [$H^3$]thymidine. (C) Cell proliferation in DAPT- (10 μM) or DMSO-treated and untreated MLE12 cells was assessed by [$H^3$]thymidine incorporation.(NT, not treated MLE12). (D)Cell proliferation in Pofut1 siRNA, scrambled (Scr) siRNA, and mock-transfected MLE12 cells, was assessed by [$H^3$]thymidine incorporation. Data in B-D are given as the mean ± SEM of the percentage of proliferating cells. *p < 0.05, **p < 0.01, ***p < 0.001 by Student's t-test.

**Fig. S5.** Transgene expression in NICD1 overexpressing mice. (A) Western blot analysis of Cre recombinase and EGFP in control and in NICD1 transgenic mice treated with doxycycline for 2 or 4 weeks. Asterisks indicate animals with successful induction of both Cre recombinase and EGFP, which were chosen for further analysis. (B) Immunofluorescence analysis of EGFP (green), Abca3 (red), and Hes1 (magenta) in control mice (n = 3; upper panel, no dox) and in mice treated with doxycycline for 2 weeks (n = 3, Dox, lower panel).

**Fig. S6.** In Vitro Notch Inhibition Restores Differentiation of AEC2s. (A) Flow cytometry histogram overlay of Lysotracker uptake in MLE12 cells treated for 3 days with increasing concentrations of DAPT (0.5 $\mu$M, 1 $\mu$M, 5 $\mu$M, 10 $\mu$M). (B) Western blot analysis of NICD1 and Hes1 expression in MLE12 cells treated as in (A). β-Tubulin served as the loading control. Right, densitometric quantification of the western blots on the left. (C) Overlaid flow cytometry histograms of Lysotracker Green uptake in MLE12 cells treated with DMSO (blue) or 10 μM DAPT (orange) for 3 days. Unstained cells served as the negative control (black, all left panel). Quantification of the log [mean fluorescence intensity] (middle panel) and the percentage (frequency of parent, right panel) of the population positive for Lysotracker Green (Lysotracker Green$^{pos)}$ in DMSO- and DAPT-treated MLE12 cells (n = 3 independent replicates per condition). Data are given as mean ± SEM of log transformed densitometric values. (D) Western blot analysis of NICD1 and Hes1 expression in human IPF PCLSs treated for 4 days with increasing concentrations of DAPT (1 $\mu$M, 10 $\mu$M, 20 $\mu$M, 50 $\mu$M). β-Tubulin served as the loading control (left panel). Right, densitometric quantification of the western blots on the left. (E) Flow cytometry analysis of dissociated cells from human IPF PCLSs showing the proportion of cells in the starting population (Day 0) or from the day 4 PCLSs treated with either DMSO or DAPT. Epithelial cells were defined as DAPI$^{neg}$, CD45$^{neg}$, CD31$^{neg}$, EpCAM$^{pos}$. AEC2 were defined as DAPI$^{neg}$, CD45$^{neg}$, CD31$^{neg}$, EpCAM$^{pos}$, Lysotracker$^{pos}$. Data are given as mean ± SEM of frequency of parent. N.s. = not significant, as assessed by Student´s t-test. (F) Gating strategy for the flow cytometry identification of epithelial cells in general and AEC2 in particular, as shown in Figure 6. (G) Immunofluorescence analysis of ABCA3 (green) and mature SP-C (red) in human IPF PCLSs treated with 50 μM DAPT (right panel) or DMSO control (left panel) for 4 days. Scale bars, 50 μm
