## Supplemental Methods for "Restored alveolar epithelial differentiation and reversed human lung fibrosis upon Notch inhibition"

*Supplementary materials and methods*

*Patient and Control Groups*

The study protocol was approved by the Ethics Committee of the Justus-Liebig-University School of Medicine (No. 31/93, 29/01, and 111/08). The data and biomaterials used in this study were provided by the UGMLC Giessen Biobank and the European IPF Registry / Biobank and informed consent was obtained in written form from each subject (see overview in Supplemental Table 1). Explanted lungs (n=43 $IPF_{LTX}$, n=9 $COPD_{LTX}$) and human donor lung lobes or pieces resected for size incompatibilities (n=30; $Donor_{LTX}$) were obtained from the Dept. of Thoracic Surgery in Giessen, Germany and Vienna, Austria (Prof. Dr. Walter Klepetko), respectively. Subpleural lung tissue samples were obtained from five additional sporadic IPF ($IPF_{VATS}$) patients who underwent video-assisted thoracic surgery (VATS) for diagnostic purposes. Likewise, bronchoalveolar lavage fluid (BALF) was collected from 10 IPF patients ($IPF_{BALF}$) at the time of first diagnosis as well as from eight healthy volunteers ($HV_{BALF}$). Blood from 25 additional patients with sporadic IPF ($IPF_{Blood}$) and 50 healthy volunteers ($HV_{Blood}$) was collected for genotyping. All IPF diagnoses were made according to the American Thoracic Society (ATS)/European Respiratory Society (ERS) consensus criteria (Raghu et al, 2011), and a usual interstitial pneumonia (UIP) pattern was shown to be present in all $IPF_{LTX}$ and $IPF_{VATS}$ patients. $IPF_{BALF}$ and $IPF_{VATS}$ patients were untreated at the time of sampling.

*Transgenic Notch 1 overexpressing mice*

Triple-transgenic mice with conditional overexpression of the Notch 1 intracellular domain (NICD1) in AEC2s were obtained by crossbreeding floxed Rosa26-NICD mice (Gt(ROSA)26Sor$^{tm1(Notch1)Dam}$/J, C57BL/6J background, JAX Mice strain No. 008159) with SPC-rtTA (Tg(SFTPC-rtTA)2Jaw/J, FVB/N background, JAX Mice strain No. 016146), and tetO-Cre (Tg(tetO-cre)1Jaw/J, FVB/N background, JAX Mice strain No 006224). Rosa26–NICD1 mice were obtained from The Jackson Laboratory, and the activator lines SP-C rtTA and tetO-Cre were kind gifts from J. Whitsett, Children's Hospital, Cincinnati, OH. Mice were crossed for at least six generations and kept in a mixed background. In resulting SPC-rtTA × tetO-Cre × NICD fl/fl triple-transgenic mice in the absence of Cre recombinase, transcription of the Rosa-NICD1 locus is blocked by a STOP sequence. Conditional Cre recombination and excision of flanking loxP sites with deletion of the STOP sequence was achieved by feeding a doxycycline-enriched standard lab chow (650 ppm doxycycline; Altromin). Littermates without transgene induction served as controls.

*Bleomycin Model of Lung Fibrosis and Chronic Pepstatin Administration*

C57BL/6N mice (Charles River Laboratories, Sulzfeld, Germany) weighting 18–22 g were used in all experiments. Mice were anaesthetized with isoflurane (Baxter) and orally intubated under a binocular loupe. Bleomycin (Hexal) in 0.9% sodium chloride (saline) at a dose of 5 U/kg body weight was administered as an aerosol using a Microsprayer (Penncentury). The specific napsin A inhibitor pepstatin A (Applichem, Darmstadt, Germany; 75 mg/kg body weight in 200 µL of a saline solution containing 10% ethanol) was administered daily over 4 weeks as an intratracheal bolus. In both experiments controls received the solvent only. At indicated time points mice were sacrificed and their lungs harvested for further analysis.

*Isolation of AEC2s and cell culture*

For AEC2 isolation, mouse lungs were perfused with 10 ml saline until visually cleared of blood, and dispase (BD Bioscience) was instilled into the lungs followed by 0.5 ml of 1% low-melting-point agarose (Sigma-Aldrich) in DMEM (Gibco). Agarose solution was allowed to solidify for a few minutes, and the lungs were then separated from the trachea and other connective tissues. The isolated organ was incubated in 2 ml of dispase for 45 min at room temperature. The lungs were then dissected in 7 ml of DMEM with DNase I (10 µg/ml; Sigma-Aldrich), minced, and dissociated by repeated pipetting into additional DMEM with DNase I. The resulting cell mixture was filtered through 70-µm, 40-µm, and 10-µm Nitex filters (BD

Bioscience). Cells were counted with Trypan Blue (Sigma-Aldrich) and then resuspended in 5 ml of DMEM supplemented with 10% FBS in the presence of the following biotinylated antibodies: anti-CD16/32, anti-CD45, and anti-CD31 (see Table S5). After incubation at 37°C for 30 min, cells were centrifuged and resuspended in DMEM containing streptavidin-coated magnetic beads (Invitrogen). The cells were incubated at room temperature for 30 min and then placed on a magnetic separator for another 15 min. The cell suspension was carefully aspirated from the beads, transferred to a new tube, centrifuged, and resuspended in DMEM. The AEC2s were plated in 48-well plates (150,000 cells/well) or used for cytospin preparations. Cells were grown up to 72 h in DMEM supplemented with 1% FBS, 1% penicillin/streptomycin, and 10 mM HEPES. All cultures were maintained in a humidified atmosphere with 5% $CO_2$ at 37°C.

*Microarrays*

*Human microdissected alveolar septa:* Laser microdissection of non-transplanted donor and explanted patient lungs was performed as described [57]. Cryosections from lung tissue were mounted on glass slides and subjected to hemalaun staining (Waldeck-Chroma). Septa were microdissected under optical control using the Laser Microbeam System (P.A.L.M.). Total RNA was isolated using the RNeasy kit (Qiagen) and further processed using the BD Atlas SMART Fluorescent Probe Amplification Kit (Clontech Laboratories, Heidelberg, Germany) with 22 SMART PCR cycles and four additional PCR cycles in the presence of aminoallylated UTP. The coupled monofunctional reactive Cy3 and Cy5 dyes were obtained from Amersham (Freiburg, Germany). The labeled dsDNA was purified with the QIAquick PCR Purification Kit (Qiagen). Labeled dsDNA containing 40 pmol incorporated Cy3 or Cy5 dyes for each sample was subjected to hybridization. IPF and donor samples were hybridized competitively on Agilent whole human genome arrays (G4112A) and washed according to Agilent's protocol (QuickAmp labeling kit, version 4.1). Slides were scanned with the Axon 4100A (Molecular Devices, Munich, Germany). Photomultiplier tube gains were adjusted to use the entire dynamic range of the scanner, yielding similar intensity histograms for both dyes.

*MLE12 cells transfected with Notch1 Intracellular Domain* (*NICD1*): To investigate the influence of Notch activation on global gene expression, MLE12 cells were transiently transfected with the NICD1-pIRES-dsRed2 vector or the pIRES-dsRd2 vector (i.e., empty vector) for 12, 24, and 48 h using Lipofectamine 2000 (Invitrogen) according to the manufacturer's instructions. Total RNA was isolated using the RNeasy kit (Qiagen) and further processed with the Agilent QuickAmp labeling kit (version 5.7). The labeled amplified RNA from NICD1 and empty vector–transfected cells was hybridized competitively according to the Agilent protocol on Agilent Whole Mouse Genome Microarrays (G4122F). Slides were scanned with the Axon 4100A. Photomultiplier tube gains were adjusted to use the entire dynamic range of the scanner, yielding similar intensity histograms for both dyes.

*Cloning of the Mouse NICD1*

To subclone the intracellular domain of Notch1 into an expression vector, the cDNA sequence was amplified by using the forward and reverse primers 5´-CGT GGC TCC ATT GTC TAC CT-3´ and 5´-CAC ACA GGG AAC TTC ACC CT-3´, respectively. The purified PCR product was ligated into pGEM-T Easy vector (Promega) and the ligation mixture was transformed into competent *E. coli* TOP10 cells (Invitrogen). To subclone NICD1 from the pGEM-T Easy vector into the mammalian expression vector pIRES-DsRed2, both the empty expression vector and pGEM-T Easy–NICD1 were digested with restriction enzymes for 1–3 h at 37°C, separated by agarose gel electrophoresis, gel-purified, and ligated. The NICD1-pIRES-DsRed2 construct was verified by sequencing.

*Mutation Analysis of the SP-C (SFTPC), SP-B (SFTPB), and napsin A (NAPSA) Genes*

Genomic DNA was extracted from fresh frozen lung tissue or EDTA whole blood using the DNeasy Blood & Tissue Kit (Qiagen). Briefly, the coding exons of *SFTPC*, *SFTPB*, and *NAPSA*, including the 5´-flanking promoter regions, were amplified with the use of primers listed in the Table S8 following a published approach [58]. Then, we digested the PCR products with shrimp alkaline phosphatase and exonuclease I and performed cycle sequencing using BigDye terminator mix (Applied Biosystems) and the oligonucleotides listed in the Table S8. The reaction products were purified by ethanol precipitation and loaded onto an ABI 3730 sequencer (Applied Biosystems). We sequenced *SFTPC*, *SFTPB*, and *NAPSA* in both directions.

*Compliance Measurements in Mice*

Lung compliance was measured in a small-animal whole-body plethysmographic chamber (Robertson Box). Mice were tracheotomized and ventilated in a volume-driven mode at a positive end-expiratory pressure (PEEP) of 0 mm Hg. Respiration rate was set at 20 breaths/min and ventilation pressure was recorded while inflating the lung at a tidal volume of 200 µL. Ventilator compliance was obtained in milliliters per kilopascals and was further corrected for animal weight.

*Harvesting Lung Tissue and Bronchoalveolar Lavage (BAL)*

After sacrifice, the thorax was opened and the lung vasculature perfused with saline solution. For BAL, the left main bronchus was ligated and a tracheotomy was performed. The right lung was lavaged by instillation of three 350-$\mu$l volumes of saline. The recovered BAL fluid was pooled and centrifuged at 300×g for 5 min to remove cells and debris, and the supernatant was immediately frozen at –80°C. The main right bronchus was ligated and cut distally from the clamp. The right lung was excised and shock-frozen in liquid nitrogen for further analysis. Next, the main left bronchus was re-opened and the left lung was filled and fixed with phosphate-buffered saline (PBS)-buffered 4% formaldehyde solution at a constant hydrostatic pressure of 20 cm $H_2O$. Next the trachea was ligated and the lung was carefully removed and transferred into the 4% formaldehyde solution. After overnight incubation at 4°C, the left lung was stored at 4°C in PBS. For histology, this lung tissue was dehydrated in a vacuum dryer and then embedded in paraffin.

*Preparation of Lung Homogenates for SDS-PAGE and Western Blot Analysis*

Peripheral human lung tissue (size 1 $cm^3$) or mouse lung tissue samples (size 0.5 $cm^3$) were pulverized in liquid nitrogen by using a mortar and pestle. The resulting powder was then transferred to 1 ml (mouse lung) or 3 ml (human lung) lysis buffer (50 mM Tris-HCl, pH 7.5; 150 mM NaCl; 1% (w/v) Triton X-100; 0.5% (w/v) Na-deoxycholate; 5 mM EDTA; and 2 mM PMSF) and incubated on ice for 1 h. Alternatively, the tissue was homogenized in lysis buffer using the Precellys tissue homogenizer (Bertin Corp.). Cell debris was removed from crude extracts by centrifugation at 10000×g for 10 min. The resulting lung homogenates were divided into aliquots and frozen at −80°C until used. Protein concentrations were determined with the Pierce BCA protein assay kit (Thermo Fisher Scientific). Lung homogenates were then diluted (1:3) in 4× SDS sample buffer (final concentration: 2% (w/v) SDS; 2.5% (v/v) β-mercaptoethanol; 10% (v/v) glycerol; 12.5 mM Tris-HCl, pH 6.8; and 0.1% (w/v) bromophenol blue) and denatured by heating at 99°C for 15 min.

Denatured lung proteins from each sample (10–50 µg/lane) were then separated by 10%, 12%, or 15% Laemmli SDS-PAGE. Thereafter, the separated proteins were transferred to a PVDF membrane (GE Healthcare) in a semi-dry blotting chamber according to the manufacturer's protocol (Bio-Rad, Munich, Germany). Immunoblots were then blocked by incubating at room temperature for 1 h in blocking buffer, 1× Tris-buffered saline (TBS; 50 mM Tris-HCl, pH 7.5; 50 mM NaCl) containing 5% (w/v) nonfat dried milk or 5% (w/v) bovine serum albumin (BSA) and 0.1% (w/v) Tween 20, followed by immunostaining for proteins of interest. Blots were incubated with the respective primary antibody (see Table S5) overnight at 4°C with gentle shaking. The blots were then washed four times in 1× TBS containing 0.1% (w/v) Tween 20 and incubated with their respective horseradish peroxidase–conjugated secondary antibodies (Table S5) for 2 h at room temperature. After four washes, blot membranes were developed with the ECL Plus chemiluminescent detection system (GE Healthcare Life Sciences) or with the Immobilon Western Chemiluminescent HRP Substrate (Millipore), and emitted signals were detected with chemiluminescent-sensitive films (Amersham Hyperfilm ECL, GE Healthcare) or with a chemiluminescence imager (Intas ChemoStar, Intas). Thereafter, blots were stripped using stripping buffer (2% (w/v) SDS and 50 mM dithiothreitol in TBS) under gentle shaking at 60°C for 1 h, followed by re-probing the blots using an antibody against the loading control protein β-actin, GAPDH, or β-tubulin. For quantification, band intensities were analyzed by densitometric scanning and were quantified using ImageJ software (NIH, version 1.46r) and normalized to the loading control.

*Immunofluorescence and Immunohistochemistry*

For immunofluorescence and immunohistochemistry (IHC) staining, standard protocols were applied. Briefly, sections (3 $\mu$m) were cut from paraffin-embedded mouse lungs and mounted on positively charged glass slides (Super Frost Plus, Langenbrinck). Sections were deparaffinized in xylene and rehydrated in graded alcohol. With the exception of immunofluorescence staining for mature SP-B and mature SP-C, heat-induced antigen retrieval in 10 mM trisodium citrate buffer (pH 6.0) was performed. Lung sections were washed for 2 min in PBS, followed by blocking in PBS containing 5% (w/v) BSA, 2% (v/v) normal donkey serum (Jackson ImmunoResearch), and 0.1% Triton X-100 (Sigma) for 30 min. For immunofluorescence staining, sections were incubated overnight at 4°C with the appropriate antibodies (Table S5). Lung sections were then washed three times in PBS (5 min), followed by incubation with the appropriate fluorochrome-conjugated secondary antibodies (Table S5). Slides were washed one time in PBS (5 min) and were then incubated with Sudan Black (Sigma-Aldrich; 3% (w/v) in 70% ethanol) for 2 min. After extensive washing in PBS, slides were incubated with 4′,6-diamidino-2-phenylindole dihydrochloride (DAPI; Sigma-Aldrich) and mounted in Fluorescence Mounting Medium (Dako). Alternatively, slides were mounted in Vectashield containing DAPI (Vector Laboratories, Burlingame). As a negative control, the first antibody was omitted and the lung tissue slides were incubated only with fluorochrome-conjugated secondary antibodies.

Two particular antibodies required signal amplification. Hes1 (clone D6P2U Cell Signaling Technologies) was used at a dilution of 1:2000 and the signal was amplified with the Alexa Fluor™ 555 Tyramide SuperBoost™ Kit, goat anti-rabbit IgG (Thermo Scientific). Cleaved Notch1 (Val1744) ((D3B8) Cell Signaling Technologies) was used at a dilution of 1:200 and the signal was amplified with the Biotin XX Tyramide SuperBoost™ Kit, Streptavidin (Thermo Scientific) following manufacturer's instructions.

Slides were imaged on a fluorescence microscope (Axio Observer.Z1 fluorescence microscope, Carl Zeiss MicroImaging, Germany). All images were acquired using the Axio Observer ZEN software, version 1.0 (Carl Zeiss MicroImaging).

Collagen 1 immunofluorescence quantification was performed using ImageJ software. Briefly, eight to ten fields (10x magnification) were acquired per PCLS (n=3 PCLS per patient, n=5 patients) and mean fluorescence intensity density (IntDen) was calculated following image thresholding and background subtraction.

For IHC we used the streptavidin–biotin–alkaline phosphatase (AP) method with the ZytoChem-Plus AP Kit (Fast Red; Zytomed Systems). Counterstaining was performed with Hemalaun (Mayer's hemalaun solution; WALDECK Division CHROMA GmbH & CO KG) for 2 min. Histological stains, Haematoxilin & Eosin and Masson's Trichrom were performed according to standard protocols. Sections were then mounted in Glycergel Mounting Medium (DakoCytomation) and imaged using a Mirax Desk scanning device (Carl Zeiss Microscopy GmbH).

*Quantitative Real-Time PCR (qPCR)*

Total RNA was extracted from human or murine lung tissue using the RNeasy® Plus Mini-Kit (Qiagen, Germany). RNA (1 $\mu$g) was then reverse-transcribed using Omniscript-RT-Kit

(Qiagen) and oligo-dT primers. qPCR was performed using SYBR Green (iQ SYBR Green Supermix Kit; Bio-Rad) and gene-specific primers (Table S8) and read on the iQ5 detection system (Bio-Rad, Germany). *ACTB*/*Actb* was used as the reference gene. Relative gene expression is presented as the ΔCt value (ΔCt = Ct reference gene – Ct gene of interest).

*Semiquantitative Reverse Transcription–PCR (RT-PCR)*

Total cellular RNA was prepared from frozen human and mouse lung tissue by an acid phenol extraction method (Korfei et al., 2008) and quantified by spectrophotometry at 260 nm.

RT and PCR were performed sequentially. cDNA (2 μg) was first synthesized by RT using 2 μg total RNA and oligo-dT primers (Applied Biosystems) with the Omniscript RT Kit (Qiagen, Hilden, Germany). An aliquot of the resulting cDNA (100 ng) was then used for PCR amplification using gene-specific primers (Table S8). For amplification of all described mRNAs, an optimized cycling protocol according to the manufacturer's protocol was used: an initial activation step of 95°C for 20 min (i.e., hot start) was followed by 25–30 three-step amplification cycles (denaturation at 94°C for 30 s, annealing at 55–60°C for 30 s, and extension at 72°C for 1 min), with a 10-min extension for the final cycle. The thermal cycler used was from Perkin Elmer (GeneAmp PCR System 2400). No reverse transcriptase controls were performed and processed in parallel.

Equal aliquots of the PCR products were separated on a 2% (w/v) agarose gel containing ethidium bromide in 1× Tris-acetate-EDTA (TAE) buffer and documented by scanning using a UV imager (AlphaEase®FC Imaging System, FluorChem™ IS 8900 software, version 3.2.3.; San Leandro, CA). Thereafter, band intensities of PCR products were quantified using ImageJ software, and mRNA expression of genes of interest was normalized to the expression of *ACTB*/*Actb*.

*Determination of Phospholipid (PL) Content and Surfactant Lipids Phosphatidylcholine (PC) and Phosphatidylglycerol (PG) in Human Lung Tissue*

Lipids were extracted from 25-μl samples of lung homogenates with 2:1 (v/v) methanol/chloroform as described (Bligh & Dyer), and PL content was determined by spectrophotometric measurement of phosphorus as published by our group [30]. High-performance thin-layer chromatography (HPTLC) was then used for the quantitative analysis of PL subclasses, as described [30], by using silica 60 plates (Merck) and 50:37.5:3.5:2 (v/v/v/v) $CHCl_3/CH_3OH/CH_3COOH/H_2O$ as the developing solvent. This one-dimensional system allows the separation of eight PL classes: lyso-phosphatidylcholine (L-PC), sphingomyelin (SPH), PC, phosphatidylserine (PS), phosphatidylinositol (PI), phosphatidylethanolamine (PE), PG, and cardiolipin (CL). Samples (25 μg total PL) were dissolved in 60 μl 2:1 (v/v) $CHCl_3/CH_3OH$ and applied together with seven concentrations of each PL standard (all from Sigma) to the plates with a Linomat IV applicator (Camag, Muttenz, Switzerland) and stained with molybdenum blue reagent [30]. Quantification was performed by densitometric scanning at 700 nm by using the thin-layer chromatography scanner II (Camag). The content of PC and PG is shown as a percentage of all PLs.

*Biophysical Properties of Alveolar Surfactant*

BAL-derived large surfactant aggregates (LSAs) were isolated from 15 IPF patients and 5 healthy volunteers by high-speed centrifugation (48,000×g, 1 h, 4°C, Sorvall centrifuge) and pooled (IPF, Control). Surface activity of the LSA pools was measured 5 times at a constant PL concentration of 2 mg/ml by means of a pulsating bubble surfactometer as described (Gunther et al, 1996). The adsorption properties after 12 sec of film adsorption (γads) and the minimum surface tension after 5 min of film oscillation ( γmin) are given. For reconstitution experiments of the IPF LSA pool, bovine SP-B was isolated from a bovine calf lung lavage extract by LH-60 chromatography. Required amounts of isolated SP-B (0.5–8% of total PL) were dried under nitrogen and resuspended with LSA fractions (PL = 2 mg/ml) by brief sonication for 30 sec (50 W, 25 kHz) and incubated for 30 min at 37°C.

*ProSP-B Electrochemiluminescence Immunoassay (ECLIA)*

For quantification of ProSP-B in serum, an ECLIA, as recently developed by Roche Diagnostics, was used. The ProSP-B assay uses mouse monoclonal anti-proSP-B (N terminus) antibodies for capture and mouse monoclonal anti-proSP-B (C terminus) antibodies as a detection reagent in a sandwich format, with a Tris(bipyridyl)-ruthenium(II) complex as the label. First, the biotinylated capture antibody, the ruthenium-labeled detection antibody, and the sample or standard material (10 $\mu$l) were incubated in homogeneous phase for 9 min at 37°C. Streptavidin-coated beads were then added. Binding of the formed immune complexes to the microparticles took place during a second 9-min incubation. After the second incubation, the reaction mixture was transferred into the measuring cell, where beads were captured on the electrode surface by a magnet. The measuring cell was washed to remove unbound label and filled with detection buffer containing Tris-propylamine. After applying voltage to the electrode, the emitted chemiluminescence light was detected by a photomultiplier. Results were determined via a two-point calibration curve. ProSP-B data are shown with the units picograms per milliliter.

*Pharmacologic and Genetic Manipulation of MLE12 Cells*

For siRNA experiments, murine alveolar MLE12 cells were cultured to 80% confluency. *Napsa* [NM_008437]-specific siRNA (sense: 5´-GGA CCA AGU UUG CCA UUC AUU-3´; antisense: 5´-P-UGA AUG GCA AAC UUG GUC CUU-3´) and non-targeting siRNA (sense: 5´-UAG CGA CUA AAC ACA UCA AUU-3´; antisense: 5´-P-UUG AUG UGU UUA GUC GCU AUU-3´) were purchased from Dharmacon Inc. MLE12 cells in 6-well plates were transfected with 100 nM *Napsa*-specific siRNA by using DharmaFECT™ 1 (Dharmacon) as the transfection reagent (cationic lipid-mediated transfection) according to the manufacturer's protocol. Untreated cells and cells transfected with non-targeting siRNA were used as negative controls. After 2 days in culture, total RNA was isolated from cells and then analyzed for gene expression of *Napsa*, *Ctsh*, *Sftpb*, *Sftpc*, *Titf1*, and *Actb*, full length and spliced *Xbp1* splicing by RT-PCR (see above). For protein analysis by immunoblotting, cells were harvested after 6 days in culture. The complete list of primers used is given in Table S8.

For *Pofut1* downregulation, siRNA from DharmaFECT™(Thermo Fisher, cat no. D-059834-01 and D-001210-03-05) was transfected as described above, and cells were cultured for 54 h before proliferation was assessed.

For Notch pharmacological inhibition, MLE12 cells were incubated with DAPT (0.5–50 µM; Selleckchem) or vehicle (dimethylsulfoxide, DMSO) for up to 72 h.

*Proliferation Assay*

MLE12 cells were exposed to [$^3$H]thymidine (0.2 µCi/well; PerkinElmer) for 6–12 h, rinsed three times with PBS, and solubilized in 0.2 ml 0.5 M sodium hydroxide. Then 0.1 ml of the solubilized material was quantified by liquid scintillation counting (TRI-CARB 1500; A Canberra Company).

*Napsin A Activity Assay*

Napsin A activity was determined by cleavage of the fluorescence resonance energy transfer (FRET)-based substrate MGAS-1 (Qxl520-KKTSVLMAAPQ-Lys–HiLyte Fluor 488; AnaSpec, San Jose, CA, USA;[59] in a microtiter plate–based assay. In brief, a 50-$\mu$l sample (lung tissue homogenate or cell lysate with equal protein amounts in saline containing 1 mM PMSF) was transferred to a 96-well microtiter plate, mixed with 140 $\mu$l of reaction buffer (0.1 M sodium acetate, 2 mM EDTA, 1 mM PMSF, and 5 mM E-64, pH 4.7), and the reaction was initiated by addition of 10 $\mu$l of the substrate**,** MGAS-1 (final concentration, 10 $\mu$M). The increase in fluorescence using excitation at 485 nm and emission at 535 nm was recorded as a function of time. Napsin A activity is expressed as fluorescence intensity units (RFU) per minute and was calculated from the slope during the linear phase of the cleavage.

*High-Precision Cut Lung Slices (PCLS) Generation and Culture*

One segment of each explanted human IPF lung was filled with 1.5% Low Melt Agarose (Bio-Rad) at 37°C and allowed to cool on ice for 30 min. Blocks of tissue of ~3 × 3 × 4 cm (depth × width × height) were cut and prepared for sectioning using a Vibrating Blade Microtome (Thermo Fisher) at a thickness of 500 µm. PCLSs were cultured for 4 days in DMEM/F12 (Gibco) supplemented with 10% human serum (Human Serum Off The Clot, Biochrom) and 1% penicillin/streptomycin in the presence of DAPT (10–50 µM in DMSO) or DMSO (1:1000; Sigma-Aldrich), during which some of the PCLSs were subjected to live imaging (see below). The medium was changed daily. At the end of the experiment, PCLSs were (a) fixed with 4% PFA for 20 min at room temperature, embedded in paraffin, and sectioned at 3 µm for immunofluorescence staining; (b) homogenized in the lysis buffer used for western blot analysis (see above); or (c) dissociated using dispase (1:10 dilution; Roche) and 10 µg/µl DNase I for 45 min at 37°C and processed for flow cytometry (see below).

*Live Imaging of Human PCLSs*

PCLSs were cultured for 4 days in the continuous presence of Lysotracker Green (Thermo Fisher Scientific), and the culture medium was changed every day. To limit movement during imaging, PCLSs were placed on top of a thin layer of Growth Factor Reduced Matrigel (BD Biosciences) and allowed to polymerize for 30 min at 37°C. Three fields per PCLS were imaged using a 10× objective every hour for 4 days using a Leica DMI6000 B live imaging microscope (Leica Microsystems) equipped with Leica DFC365 FX camera. Images were acquired and processed using the Leica Application Suite X software (Leica Microsystems).

*Flow Cytometry*

Dissociated cells from PCLSs cultured as described above were counted and resuspended at $10^6$ cells/100 µl and incubated with the appropriate directly conjugated antibodies or Lysotracker Green (100 nM; Table S5 and S6) in fluorescence-activated cell sorting (FACS) buffer, which consists of Hanks buffered salt solution (HBSS; Pan Biotech) supplemented with 2% FBS (PAA), 0.1 mM HEPES, 2 µg/µl DNase I (Sigma-Aldrich), and 1% penicillin/streptomycin (Gibco), for 30 min at 37°C. For the MLE12 analysis, MLE12 cells were trypsinized and resuspended at $10^6$ cells/100 µl in FACS buffer containing 100 nM Lysotracker Green and were incubated for 30 min at 37°C. Following incubation, samples were washed one time with 1 ml FACS buffer and resuspended in 200 µl FACS buffer with DAPI.

Human lung cell suspension was thawed rapidly and cells were washed in 10 times the freezing volume of RPMI 10% FBS and maintained on ice throughout the procedure. Cells were resuspended in FACS buffer at a concentration of $10^6$ cells/100 µl and cell surface staining was performed as described for PCLS. For intracellular staining, cell surface was first performed, then cells were fixed and permeabilized for 30 min at room temperature using the Foxp3 / Transcription Factor Staining Buffer Set (eBioscience – Thermo Scientific) according to the manufacturer instructions. Cells were washed following fixation and resuspended in 100ul of Permeabilization/wash buffer containing mature SP-B antibody diluted 1:200, incubated for 1hour at room temperature followed by Anti-rabbit IgG (H+L), $F(ab')_2$ Fragment (Alexa Fluor® 555 Conjugate Cell Signaling Technologies 1:1000). Data were acquired on a BD FACSCanto II (BD Biosciences) using BD FACSDiva software (BD Biosciences). Data were further analyzed using FlowJoX software (FlowJo, LLC).

### Quantification and statistical analysis

*Microarray Data Analysis*

Scanned microarray images were analyzed using GenePix 5.0 (Molecular Devices), and further data processing was performed using R and the limma package [60] (Smyth, 2004). Data from laser-microdissected human samples were lowess normalized, data from cell culture experiments were quantile normalized. Genes were ranked for differential expression using a moderated t-statistic (Smyth, 2004). Pathway analyses were done by gene set tests using the ranks of the t-statistics and the pathway annotation from KEGG [61].

*Quantitative Lung Morphometry*

For morphometry analysis of Haematoxylin & Eosin (H&E)-stained lung sections, the entire section was scanned with a Mirax Desk scanning device (Carl Zeiss Microscopy GmbH, Jena, Germany), and the resulting digital image was subjected to further analysis with the Axiovision® Software plug-in (Carl Zeiss Microscopy GmbH, Jena, Germany). The method consisted of generating a graphic overlay of parallel horizontal and vertical chords over representative areas of the lung section (chord distance, 20 $\mu$m). For each lung slide, ~20–25 representative fields covering the subpleural area of the lung were defined and subjected to analysis. Almost the entire peripheral lung area of each section was subjected to analysis with this approach, although hilar structures like bronchi and large vessels were excluded. The number and length of each chord for septal thickness and air space, respectively, was calculated, resulting in about 30,000–60,000 single values per individual section. The probability density function of septal thickness and alveolar mean linear intercept was determined by kernel density estimation of chord length (Parzen-window-method) [62]. A kernel is defined as the density of a Gaussian distribution $N(\mu,\sigma^2)$ with a smoothening factor (bandwidth) of 0.2 $\mu$m. The displayed curves as given in Figure 6 F represent point-wise mean values of individual densities within a group, and the shaded area indicates the standard errors.
