## Supplemental Tables for "Restored alveolar epithelial differentiation and reversed human lung fibrosis upon Notch inhibition"

**Table S1.** Basic characteristics of the patients involved in our studies

| Patient category | n | Age (mean ± SEM) | Gender (m/f) | FVC (mean ± SEM) | FEV1 (mean ± SEM) | Smoking status %current/former/ never | PY (mean ± SEM) | UIP pattern (n/n open biopsies) |
|---|---|---|---|---|---|---|---|---|
| Idiopathic Pulmonary Fibrosis | | | | | | | | |
| $IPF_{LTX}$ | 47 | 54.4 ± 1.6 | 39/18 | 43.8 ± 2.5 | 48.6 ± 2.7 | 0/59/41 | 23.2 ± 5.6 | 47/47 |
| $IPF_{VATS}$ | 5 | 61.0 ± 1.9 | 3/2 | 61.1 ± 6.7 | 66.6 ± 6.2 | 0/60/40 | 15.0 ± 7.5 | 5/5 |
| $IPF_{BALF}$ | 10 | 64.7 ± 2.5 | 6/4 | 71.8 ± 7.7 | 75.1 ± 6.0 | 10/30/60 | 25.0 ± 17.5 | 5/5 |
| $IPF_{Blood}$ | 25 | 61.9 ± 2.7 | 17/8 | 61.2 ± 4.1 | 66.3 ± 4.9 | 4/28/68 | 19.8 ± 16.3 | 9/9 |
| Chronic Obstructive Lung Disease/Emphysema | | | | | | | | |
| $COPD_{LTX}$ | 9 | 55.2 ± 1.4 | 6/3 | 38.4 ± 3.0 | 17.0 ± 1.2 | 0/9/0 | 51.8 ± 7.7 | 0/9 |
| Control Subjects | | | | | | | | |
| $Donors_{LTX}$ | 43 | 47.0 ± 2.8 | 20/23 | n.a. | n.a. | n.a. | n.a. | n.a. |
| Healthy $volunteers_{BALF}$ | 8 | 24.8 ± 1.3 | 6/2 | 103.3 ± 2.6 | 106.7 ± 2.7 | 0/0/8 | 0 | n.a. |
| Healthy $volunteers_{Blood}$ | 50 | 34.4 ± 1.4 | 27/23 | n.a. | n.a. | n.a. | n.a. | n.a. |

*Abbreviations:* BALF = bronchoalveolar lavage fluid; FEV1 = forced expiratory volume after 1 sec; FVC = forced vital capacity; LTX = lung transplant; PY = pack years; n.a. = not applicable.

**Table S2.** *SFTPB*, *SFTPC*, and *NAPSA* Genotyping in IPF patients and healthy controls

| Gene | Polymorphism | [Allele type[a]] | Controls (n = 50) | IPF (n = 36) (11 $IPF_{LTX}$ + 25 $IPF_{Blood}$) |
|---|---|---|---|---|
| *SFTPB* | I131T | wt | 11/50 (22%) | 18/36 (50%) |
| | | heterozygous | 28/50 (46%) | 15/36 (42%) |
| | | homozygous | 11/50 (22%) | 3/36 (8%) |
| | I102V | wt | 50/50 (100%) | 35/36 (97%) |
| | | heterozygous | 0/50 (0%) | 1/36 (3%) |
| | | homozygous | 0/50 (0%) | 0/36 (0%) |
| | E97E(gAg>gAA) | wt | 50/50 (100%) | 35/36 (97%) |
| | | heterozygous | 0/50 (0%) | 1/36 (3%) |
| | | homozygous | 0/50 (0%) | 0/36 (0%) |
| | T16T(ACg>ACA) | wt | 50/50 (100%) | 35/36 (97%) |
| | | heterozygous | 0/50 (0%) | 1/36 (3%) |
| | | homozygous | 0/50 (0%) | 0/36 (0%) |
| *SFTPC* | T138N | wt | 25/50 (50%) | 20/36 (56%) |
| | | heterozygous | 21/50 (42%) | 16/36 (44%) |
| | | homozygous | 4/50 (8%) | 0/36 (0%) |
| | S186N | wt | 21/50 (42%) | 15/36 (42%) |
| | | heterozygous | 24/50 (48%) | 16/36 (44%) |
| | | homozygous | 5/50 (10%) | 5/36 (14%) |
| *NAPSA* | I40T | wt | 24/50 (48%) | 12/36 (33%) |
| | | heterozygous | 21/50 (42%) | 19/36 (53%) |
| | | homozygous | 5/50 (10%) | 5/36 (14%) |
| | P255L | wt | 5/50 (10%) | 5/36 (14%) |
| | | heterozygous | 21/50 (42%) | 19/36 (53%) |
| | | homozygous | 24/50 (48%) | 12/36 (33%) |
| | A310T | wt | 41/50 (82%) | 33/36 (92%) |
| | | heterozygous | 9/50 (18%) | 3/36 (8%) |
| | | homozygous | 0/50 (0%) | 0/36 (0%) |

[a]For the allele type data, the frequency of two wild-type (wt) alleles, of a mutant and wild-type allele (heterozygous), and of two mutant alleles (homozygous) is shown.

**Table S3.** Top ten signaling pathways that are differentially regulated in the normal appearing septa from IPF patients versus donor septa and in fibrotic septa from IPF patients versus donor septa

| Normal Appearing Septa vs Donor Septa | | Fibrotic Septa vs Donor Septa | |
|---|---|---|---|
| Pathway | [log p OR $log_{10}$ p][a] | Pathway | [log p OR $log_{10}$ p][a] |
| Neuroactive ligand-receptor interaction | 18.39 | AMPK signaling pathway | 2.45 |
| Cytokine-cytokine receptor interaction | 4.31 | TNF signaling pathway | 1.94 |
| Jak-STAT signaling pathway | 4.05 | Wnt signaling pathway | 1.93 |
| ErbB signaling pathway | 2.75 | PI3K-Akt signaling pathway | 1.42 |
| Cell adhesion molecules (CAMs) | 2.40 | ErbB signaling pathway | 1.39 |
| TGF-beta signaling pathway | 2.38 | NF-kappa B signaling pathway | 1.17 |
| ABC transporters | 2.13 | TGF-beta signaling pathway | 1.17 |
| Notch signaling pathway | 2.04 | Hippo signaling pathway | 1.15 |
| cAMP signaling pathway | 1.88 | Ras signaling pathway | 1.11 |
| Ras signaling pathway | 1.69 | Cell adhesion molecules (CAMs) | 1.08 |

[a]p-values for each pathway were log transformed.

**Table S4.** Differentially regulated cellular processes in MLE12 Cells after 24 and 48 h of NICD1 upregulation

| Processes regulated only after 24 h | Processes regulated after 24 h and 48 h | Processes regulated only after 48 h |
|---|---|---|
| • Antigen processing and presentation<br>• Atrazine degradation<br>• Cell adhesion molecules (CAMs)<br>• Cytokine-cytokine receptor interaction<br>• Huntington's disease<br>• Jak-STAT signaling pathway<br>• Long-term depression<br>• Melanogenesis<br>• Natural killer cell mediated cytotoxicity<br>• T cell receptor signaling pathway | • Acute myeloid leukemia<br>• Adherens junction<br>• Apoptosis<br>• Axon guidance<br>• Bladder cancer<br>• Cell cycle<br>• Chronic myeloid leukemia<br>• Colorectal cancer<br>• ECM-receptor interaction<br>• Endometrial cancer<br>• ErbB signaling pathway<br>• Focal adhesion<br>• Gap junction<br>• Glioma<br>• GnRH signaling pathway<br>• Insulin signaling pathway<br>• MAPK signaling pathway<br>• Melanoma<br>• mTOR signaling pathway<br>• Pancreatic cancer<br>• Prostate cancer<br>• Regulation of actin cytoskeleton<br>• Small cell lung cancer<br>• TGF-beta signaling pathway<br>• Toll-like receptor signaling pathway<br>• Type II diabetes mellitus<br>• Ubiquitin mediated proteolysis | • Adipocytokine signaling pathway<br>• Alzheimer's disease<br>• Aminoacyl-tRNA biosynthesis<br>• Aminosugars metabolism<br>• B cell receptor signaling pathway<br>• Base excision repair<br>• Biosynthesis of steroids<br>• Dentatorubropallidoluysian atrophy (DRPLA)<br>• DNA replication<br>• Fatty acid elongation in mitochondria<br>• Fructose and mannose metabolism<br>• Glycan structures - biosynthesis<br>• Glycan structures - degradation<br>• Glycine, serine and threonine metabolism<br>• Glycosylphosphatidylinositol(GPI)- anchor biosynthesis<br>• Homologous recombination<br>• Inositol phosphate metabolism<br>• Long-term potentiation<br>• Lysine degradation<br>• Mismatch repair<br>• N-Glycan biosynthesis<br>• Nicotinate and nicotinamide metabolism<br>• Non-homologous end-joining<br>• Non-small cell lung cancer<br>• Nucleotide excision repair<br>• Oxidative phosphorylation<br>• p53 signaling pathway<br>• Parkinson's disease<br>• Pentose phosphate pathway<br>• Phosphatidylinositol signaling system<br>• Proteasome<br>• Purine metabolism<br>• Pyrimidine metabolism<br>• Regulation of autophagy<br>• Renal cell carcinoma<br>• Ribosome<br>• SNARE interactions in vesicular transport |

| | | |
|---|---|---|
| | | • Thyroid cancer<br>• Tight junction<br>• Valine, leucine and isoleucine degradation<br>• VEGF signaling pathway<br>• Wnt signaling pathway |

Table S5: Antibodies used in this study

| ANTIBODIES | SOURCE | IDENTIFIER |
|---|---|---|
| anti-Notch1 | Abcam | ab52627 |
| anti-Notch1 | Abcam | ab8925 |
| anti-Notch1 | Cell Signaling Technologies | CST4147 |
| anti-Notch2 | Cell Signaling Technologies | CST4530 |
| anti-Notch3 | Abcam | ab23426 |
| anti-Notch4 | Cell Signaling Technologies | CST2423 |
| anti-Hes1 | R&D Systems | AF3317 |
| anti-Hes1 | Cell Signaling Technologies | CST11988 |
| anti-DLL1 | R&D Systems | AF 5026 |
| anti-DLL3 | Cell Signaling Technologies | CST2483 |
| anti-Jagged1 | Abcam | ab7771 |
| anti-ABCA3 | Seven Hills Bioreagents | WMAB-17G524 |
| anti-proSP-C | Seven Hills Bioreagents | WRAB-9932 |
| anti-proSP-C | SantaCruz | sc-7706 |
| anti-mature SP-C | Seven Hills Bioreagents | WRAB-76694 |
| anti-mature SP-C (recombinant human) | Nycomed | N/A |
| anti-proSP-B | Millipore | AB3430 |
| anti-mature SP-B | Seven Hills Bioreagents | WRAB-48604 |
| anti-Beta-Tubulin | Sigma | T0198 |
| anti-Beta-Actin | Abcam | ab8227 |
| anti-Beta-Actin | Abcam | ab8226 |
| anti-PCNA | Abcam | ab18197 |
| anti-PCNA | SantaCruz | sc-56 |
| anti-phospho Histone H3 | Abcam | ab5176 |
| anti-Ki67 | Abcam | ab15580 |
| anti-Collagen I | Meridian Life Sciences | T40777R |

| | | |
|---|---|---|
| anti-Collagen I | Rockland | 600-401-103 |
| anti-Vimentin | Abcam | ab92547 |
| anti-Vimentin | SantaCruz | sc58901 |
| anti-alpha SMA | Abcam | ab119952 |
| anti-alpha SMA | Abcam | ab5694 |
| anti-Cre recombinase | Biolegend | 908001 |
| anti-GFP | Millipore | AB16901 |
| anti-GFP | Abcam | ab5450 |
| anti-EpCAM Pe-Cy7 | Biolegend | 324222 |
| anti-CD45 APC-Cy7 | Biolegend | 304016 |
| anti-CD31 APC-Cy7 | Biolegend | 303120 |
| anti-CD16/32 (biotinylated) | BD Biosciences | 553143 |
| anti-CD45 (biotinylated) | BD Biosciences | 553078 |
| anti-CD31 (biotinylated) | BD Biosciences | 553371 |
| anti-Cathepsin H | Abcam | ab7432 |
| anti-Cathepsin H | St Cruz | sc6496 |
| anti-Napsin A | Abcam | ab9868 |
| anti-anti-TTF1 | Upstate/Millipore | 07-601 |
| anti-Chop | St Cruz | sc575 |
| Donkey anti-rabbit Alexa-Fluor 488 | ThermoFisherScientific | A21206 |
| Donkey anti-rabbit Alexa-Fluor 555 | ThermoFisherScientific | A31572 |
| Donkey anti-rabbit Alexa-Fluor 647 | ThermoFisherScientific | A31573 |
| Donkey anti-mouse Alexa-Fluor 488 | ThermoFisherScientific | A21202 |
| Donkey anti-mouse Alexa-Fluor 555 | ThermoFisherScientific | A31570 |
| Donkey anti-mouse Alexa-Fluor 647 | ThermoFisherScientific | A31571 |
| Donkey anti-goat Alexa-Fluor 488 | ThermoFisherScientific | A11055 |
| Donkey anti-goat Alexa-Fluor 555 | ThermoFisherScientific | A21432 |
| Donkey anti-goat Alexa-Fluor 647 | ThermoFisherScientific | A21447 |

| | | |
|---|---|---|
| Donkey anti-chicken Alexa-Fluor 488 | ThermoFisherScientific | A11039 |
| Donkey anti-chicken Alexa-Fluor 555 | ThermoFisherScientific | A21437 |
| Anti-mouse F(ab')2 Fragment Alexa Fluor 488 | Cell SignalingTechnologies | CST4408 |
| Anti-mouse F(ab')2 Fragment Alexa Fluor 555 | Cell SignalingTechnologies | CST4409 |
| Anti-rabbit F(ab')2 Fragment Alexa Fluor 488 | Cell SignalingTechnologies | CST4412 |
| Anti-rabbit F(ab')2 Fragment Alexa Fluor 555 | Cell SignalingTechnologies | CST4413 |
| HRP-conjugated rabbit anti–mouse IgG | DakoCytomation | P0260 |
| HRP-conjugated rabbit anti–goat IgG | DakoCytomation | P0160 |
| HRP-conjugated rabbit anti–sheep IgG | DakoCytomation | P0163 |
| HRP-conjugated swine anti–rabbit IgG | DakoCytomation | P0217 |

Table S6. Chemicals, peptides and recombinant proteins used in this study

| CHEMICALS, PEPTIDES, PROTEINS | SOURCE | IDENTIFIER |
|---|---|---|
| Bleomycinsulfate, injection solution 15,000 U | Hexal | ATC code: L01DC01 |
| Pepstatin A (napsin-A inhibitor) | Applichem | A2205 |
| PMSF | Sigma-Aldrich | P7626 |
| DAPT | Selleckchem | S2215 |
| E-64 | Sigma-Aldrich | E3132 |
| Fetal bovine serum (FBS) | PAA | S0615 |
| Bovine serum albumin (BSA) | Roth | 80763 |
| Human Serum | Biochrom | S01049 |
| Donkey serum | Jackson Immuno Research | 017-000-001 |
| hydro-β-estradiole | Sigma-Aldrich | E2758 |
| hydrocortisone | Sigma-Aldrich | H0888 |
| ITS solution: insulin, transferrin Na-selenite | PAN Biotech | P07-03100 |
| L-Glutamine, 200 mM | Gibco | 25030-024 |
| Penicillin/Streptomycin | Gibco | 15140-122 |
| Dispase | BD Biosciences | 354235 |
| Dispase 1:10 dilution | Roche | 04942086001 |
| Low-melting-point agarose | Sigma-Aldrich, Biorad | 161-3111 |
| DNAse I | Sigma-Aldrich | DN25-1G |
| Na-deoxycholate | Sigma Aldrich | 30970-100G |
| DMSO | Sigma Aldrich | D-8418 |
| Triton X-100 | Sigma Aldrich | T8787 |
| Tween-20 | Sigma Aldrich | P8074 |
| EDTA, disodium salt (Titriplex III) | Merck | 108418 |
| Sodium dodecyl sulfate, SDS | Roth | CN30.3 |
| β-mercaptoethanol | Sigma-Aldrich | M6250 |

| | | |
|---|---|---|
| Acrylamide/bis-acrylamide solution, Rotiphorese Gel 30 | Roth | 3029.1 |
| ECL Plus western blotting substrate | GE Healthcare | RPN2133 |
| Immobilon western chemiluminescent HRP substrate | Millipore | WBKLS0500 |
| 4,6-Diamidino-2-phenylindole dihydrochloride (DAPI) | Sigma-Aldrich | D9542 |
| Sudan black | Sigma-Aldrich | 199664-25G |
| Fluorescence Mounting Medium | DakoCytomation | S3023 |
| Vectashield with DAPI | Vector Laboratories | H-1200 |
| Glycergel Mounting Medium | DakoCytomation | C0563 |
| Mayer's hemalaun solution | Waldeck-Chroma | 2E-038 |
| lyso-phosphatidylcholine (HPTLC standard) | Sigma-Aldrich | L4129 |
| Sphingomyelin (HPTLC standard) | Sigma-Aldrich | S0756 |
| Phosphatidylcholine (HPTLC standard) | Sigma-Aldrich | P4139 |
| Phosphatidylserine (HPTLC standard) | Sigma-Aldrich | P0474 |
| Phosphatidylethanolamine (HPTLC standard) | Sigma-Aldrich | P3511 |
| Phosphatidylglycerol (HPTLC standard) | Sigma-Aldrich | P8318 |
| Cardiolipin (HPTLC standard) | Sigma-Aldrich | C0563 |
| Lipofectamine 2000 | Invitrogen | 1668027 |
| [3H]thymidine | PerkinElmer | NET355001MC |
| Napsin A fluorogenic substrate MGAS-1 (Qxl520-KKTSVLMAAPQ-Lys–HiLyte Fluor 488) | AnaSpec | MGAS-1 |
| Lysotracker Green | Thermo Fisher Scientific | L7526 |
| Matrigel, growth factor reduced | Corning | 356231 |

Table S7. Commercial assays used in this study

| COMMERCIAL ASSAYS | SOURCE | IDENTIFIER |
|---|---|---|
| RNeasy kit (total RNA isolation) | Qiagen | 74106 |
| Omniscript-RT-Kit (reverse transcription) | Qiagen | 205113 |
| iQ SYBR Green Supermix Kit (qPCR) | Bio-Rad | 1708880 |
| Human Gene Expression 4x44K Microarray Kit | Agilent Technologies | G4112A |
| Mouse Gene Expression 4x44K Microarray Kit | Agilent Technologies | G4122F |
| Backing slides | Agilent Technologies | G2534-60012 |
| BD Atlas SMART Fluorescent Probe Amplification Kit<br><br>(microarray) | Clontech Laboratories, Heidelberg, Germany | K1861-1 |
| QIAquick PCR Purification Kit (labeled DNA purification) | Qiagen | 28104 |
| QuickAmp labeling kit (v. 5.7) | Agilent | 5190-0444 |
| Hi-RPM GE Hybridization Kit | Agilent | 5190-0404 |
| Gene Expression Wash Buffer Kit | Agilent | 5188-5327 |
| Stabilization and Drying Solution | Agilent | 5185-5979 |
| DNeasy Blood & Tissue Kit (DNA isolation) | Qiagen | 69504 |
| Big Dye Terminator Mix (Sequencing) | Applied Biosystems | 4337455 |
| Pierce BCA protein assay | Thermo Fisher Scientific | 23227 |
| ZytoChem-Plus AP Kit (Fast Red IHC staining kit) | Zytomed Systems | AP008RED |
| ProSP-B ECLIA | Roche/ This paper | N/A |

Table S 8. Oligonucleotides used in this study

| OLIGONUCLEOTIDES | SOURCE | IDENTIFIER |
|---|---|---|
| Pofut1 siRNA DharmaFECT | Thermo Fisher | Cat# 059834-01 and D-001210-03-05 |
| siRNA Napsa targeting: 5´-GGA CCA AGU UUG CCA UUC AUU-3´(sense), 5´-P-UGA AUG GCA AAC UUG GUC CUU-3´ (antisense) | Dharmacon | D-001210-01-05 |
| siRNA non-targeting: 5´- UAG CGA CUA AAC ACA UCA AUU -3´(sense), 5´- P- UUG AUG UGU UUA GUC GCU AUU-3´ (ANM_005411ntisense) | Dharmacon | D-046905-02 |
| RT-PCR Primer human SFTPA: Forward, 5´- ATC TAG ATG AGG AGC TCC AAG C-3´; Reverse, 5´- CCT CAG TCA GGC CTA CAT AGG - 3´ | | NM_005411 |
| RT-PCR Primer human SFTPB: Forward 5´- AAG TGC TTG ACG ACT ACT TCC - 3´, Reverse 5´- GCT TGG ATC CGC TTG ATC AG - 3´ | | NM_198843 |
| RT-PCR Primer human SFTPC: Forward 5´- CTC ATC GTC GTG GTG ATT GTG - 3´, Reverse 5´- CTG CAG AGA GCA TTC CAT CTG - 3´ | | NM_003018 |
| RT-PCR Primer human SFTPD: Forward 5´- CCA CAG AAC AAT GCC CAG TG - 3´, Reverse 5´- TTG CCC TGA GGT CCT ATG TTC - 3´ | | NM_003019 |
| RT-PCR Primer human NAPSA: Forward 5´- TCA CCT TCG TGC CAG TCA C - 3´, Reverse 5´- TCG AAG ACG GCC ACA TAC G - 3´ | | NM_004851 |
| RT-PCR Primer human CTSH: Forward 5´- CCA TCG CAA CCG GAA AGA TG - 3´, Reverse 5´- ACA TCA TGA AGT CCT GAG TCA C - 3´ | | NM_004390 |
| RT-PCR Primer human ATP1B1: Forward 5´- AGC CCA GAG GGA TGA CAT G - 3´, Reverse 5´- TCC TTA TCT TCA TCT CGC TTG C - 3´ | | NM_001677 |
| RT-PCR Primer human ABCA3: Forward 5´- CTT CAT CAT GCC CTC CTA TTG G - 3´, Reverse 5´- TGA TGT ATG CCC GTC CAC TG - 3´ | | NM_001089 |
| RT-PCR Primer human SCGB1A1: Forward 5´- TCC GCT TCT GCA GAG ATC TG - 3´, Reverse 5´- GTG TCC ACC AGC TTC TTC AG - 3´ | | NM_003357 |

| | | |
|---|---|---|
| RT-PCR Primer human ACTB: Forward 5´- ACC CTG AAG TAC CCC ATC G - 3´, Reverse 5´- CAG CCT GGA TAG CAA CGT AC - 3´ | | NM_001101 |
| RT-PCR Primer mouse spliced XBP1: Forward 5´- AGC TTT TAC GGG AGA AAA CTC A-3’, Reverse 5´- GCC TGC ACC TGC TGC G-3’ | | NM_013842 |
| qPCR Primer mouse Notch 1: Forward 5´- atggcttcgactgccagctcac-3´, Reverse 5´- tcggcactgttacagccctggt-3´ | | NM_008714.3 |
| qPCR Primer mouse Notch 2: Forward 5´- gggcagctgctgtcaataat-3´, Reverse 5´- tttggccgcttcataacttc-3 | | NM_010928.2 |
| qPCR Primer mouse Notch 3: Forward 5´- caggccacgtgtcttgaccgaa-3´, Reverse 5´- tgggctgctctgacattcgtcg-3´ | | NM_008716.2 |
| qPCR Primer mouse Notch 4: Forward 5´- tctggatgtggacacctgtggacc-3´, Reverse 5´- tctctgtggactagccccagtcgt-3´ | | NM_010929.2 |
| qPCR Primer mouse DLL1: Forward 5´- gccttcagcaaccccat-3´, Revere 5´- tgttgcgaggtcatcgg-3 | | NM_007865.3 |
| qPCR Primer mouse´DLL4: Forward 5´- tgcctgggaagtatcctcac-3´, Reverse 5´- tagagtccctgggagagcaa-3´ | | NM_019454.3 |
| qPCR Primer mouse Jagged 1: Forward 5´- actgggcctgacaaatacca-3´, Reverse 5´- tgaggaggtctccttgcag-3´ | This paper | NM_013822.5 |
| qPCR Primer mouse Jagged 2: Forward 5´- gcctcctcctgctgctttgtga-3´, Reverse 5´- atcaggctgctgtcaggcaggt-3´ | This paper | NM_010588.2 |
| qPCR Primer mouse HES1: Forward 5´- ctgcagcgggcgcagatgac-3´, Reverse 5´- acacgtggacaggaagcggg-3´ | This paper | NM_008235.2 |
| qPCR Primer mouse HEY1: Forward 5´- ccacgctccgccaccatgaa-3´, Reverse 5´- cggcgcttctcgatgatgcct-3´ | This paper | NM_010423.2 |
| qPCR Primer mouse HEY2: Forward 5´- tcgcgatgaagcgcccttgt-3´, Reverse 5´- tcactgagcttgtagcgtgcc-3´ | This paper | NM_013904 |

| | | |
|---|---|---|
| qPCR Primer mouse beat actin: Forward 5´-ctacagcttcaccaccacag-3´, Reverse 5´-ctcgttgccaatagtgatgac-3´ | This paper | NM_007393 |
| qPCR Primer human Notch 1: Forward 5´-atggacgtcaatgtccgc-3´, Reverse 5´-ccctggtagatgaagtcgga-3´ | This paper | NM_017617.3 |
| qPCR Primer human Notch 2: Forward 5´-catggccaatagcaatcctt-3´, Reverse 5´-tcacaacgaggtcctgcata-3´ | This paper | NM_024408.3 |
| qPCR Primer human Notch 3: Forward 5´-ccgatgtcaacgagtgtctg-3´, Reverse 5´-aatgtccacctcgcaatagg-3´ | This paper | NM_000435.2 |
| qPCR Primer human Notch 4: Forward 5´-gaccagaaagacaaggccaa-3´, Reverse 5´-aacccacgtcacacacacat-3´ | This paper | NM_004557.3 |
| qPCR Primer human DLL1: Forward 5´-gaatctgtgtggagagcttcaat-3´, Reverse 5´-gtcgactccttcagtctgcc-3´ | This paper | NM_005618.3 |
| qPCR Primer human DLL4: Forward 5´-tctgacccacagctagggag-3´, Reverse 5´-tctcgctcatcatcgaagc-3´ | This paper | NM_019074.3 |
| qPCR Primer human Jagged 1: Forward 5´-caagtgccaccgtttctaca-3´, Reverse 5´-agtcgggaggcaaattcac-3´ | This paper | NM_000214.2 |
| qPCR Primer human Jagged 2: Forward 5´-gatcccggagcaaatgg-3´, Reverse 5´-ggccacctggacaataactg-3´ | This paper | NM_145159.1 |
| qPCR Primer human beat actin: Forward 5´-acagagcctcgcctttgccg-3´, Reverse 5´-acatgccggagccgttgtcg-3 | This paper | NM_007393 |
| Primer cloning mouse Notch 1: Forward 5´- CGT GGC TCC ATT GTC TAC CT- 3´, Reverse 5´- CAC ACA GGG AAC TTC ACC CT-3´ | This paper | NM_008714.3 |

<table>
<tr><td>NAPSA Genotyping, Primers for PCR amplification of fragments:<br><br>*promoter - exon 1:* Forward 5-CTGACAGCAGCTGAAGGATG-3, Reverse 5-CTAGGGATCCTGGGTGCCAA-3<br><br>*exon 2 + 3:* Forward 5-ACTATTGTAGTGCCTTGGAG-3, Reverse 5-AGCCTCTGAGAAGCTGAGGT-3<br><br>exon *4*: Forward 5-GACCTCAGCTTCTCAGAGGC-3, Reverse 5-ATTGGCTTGGGAAGCTCCTC-3<br><br>*exon 5:* Forward 5-CTAGGAAGTTGGGGCTTGCA-3, Reverse 5-GGTCAGTGACTTCCTGAAGG-3<br><br>exon *6 + 7*: Forward 5-CCTGGCAATACCTAGGGCTG-3, Reverse 5-CTGTCAAACTGCCATCAGCC-3<br><br>*exon 8: Forward 5-GGAGCCACGGAAGGGACTG-3, Reverse 5-CAACAAACTGCCATCACAGG-3*<br><br>*exon 9*: Forward 5-GTCCTTGTGGCCGCGACACC-3, Reverse 5-AGCAACCCAGGCAGGTTCGC-3</td><td>This paper</td><td>NM_004851</td></tr>
</table>

| | | |
|---|---|---|
| NAPSA Genotyping, Primers for DNA Sequencing<br><br>*promoter - exon 1*: 5-CTGACAGCAGCTGAAGGATG-3<br><br>*promoter - exon 1*: 5-CTAGGGATCCTGGGTGCCAA-3<br><br>*exon 2 + 3:* 5-ACTATTGTAGTGCCTTGGAG-3<br><br>*exon 2 + 3*: 5-AGCCTCTGAGAAGCTGAGGT-3<br><br>*exon 4:* 5-GACCTCAGCTTCTCAGAGGC-3<br><br>*exon 4*: 5-ATTGGCTTGGGAAGCTCCTC-3<br><br>*exon 5:* 5-CTAGGAAGTTGGGGCTTGCA-3<br><br>*exon 5*: 5-GGTCAGTGACTTCCTGAAGG-3<br><br>*exon 6 + 7*: 5-CCTGGCAATACCTAGGGCTG-3<br><br>*exon 6 + 7*: 5-CTGTCAAACTGCCATCAGCC-3<br><br>*exon 8*: 5-GGAGCCACGGAAGGGACTG-3<br><br>*exon 8*: 5-CAACAAACTGCCATCACAGG-3<br><br>exon *9*: 5-GTCCTTGTGGCCGCGACACC-3<br><br>*exon 9*: 5-AGCAACCCAGGCAGGTTCGC-3 | This paper | NM_004851 |
| SFTPB Genotyping, Primers for PCR amplification of fragments:<br><br>*promoter - exon 2:* Forward 5-CCTGGGTCTGCCCTTCCAGG-3, Reverse 5-TCCTCCTGCCCATCCAGAGC-3<br><br>*exon 3 + 4:* Forward 5-CAAGTTGGGCTGGTGGGCAG-3, Reverse5-TGTGTGTGGCTCCCCCATGG-3<br><br>*exon 5 + 6:* Forward 5-CCCTAAAGTCCCCACACAGC-3, Reverse 5-TACCAGGCCTGAGCCTGAGC-3<br><br>*exon 7 + 8:* Forward 5-GTAGGTCTGAAGCTGGCTCC-3, Reverse 5-GTCTGTGCTCCATTCTGGCC-3<br><br>*exon 9 + 10*: Forward 5-GTCCTCCGTCTCCAGTGTCG-3, Reverse 5-GTTGGGCACTCAGTGACTGG-3 | This paper | NM_198843 |

| | | |
|---|---|---|
| SFTPB Genotyping, Primers for DNA Sequencing<br><br>*exon 1:* 5-ACCAGATGCCCTCCACCCTC-3<br><br>*exon 2*: 5-CCTCTCCTAGGCAGCTCCAC-3<br><br>*exon 3*: 5-CTGCATGTGCCTTGGAGTGC-3<br><br>*exon 4*: 5-TCGTGAACTCCAGCACCCTG-3<br><br>*exon 5*: 5-TCCAGTGGTCCCTGAGCCCT-3<br><br>*exon 6*: 5-GAGATCCAGAGGGCTAGAGC-3<br><br>*exon 7*: 5-CCTGCATCCCCTGGACTCTC-3<br><br>*exon 8:* 5-CTACCCTGCCACTGCATGAC-3<br><br>*exon 9*: 5-AGCCAGAGGTGTTCCGTGAG-3<br><br>*exon 10*: 5-CATCTCACCTCCTCAGGCTC-3 | This paper | NM_198843 |
| SFTPC Genotyping, Primers for PCR amplification of fragments:<br><br>*promoter - exon 1:* Forward 5'-GTTGGAACTGGTCCTTGCAGG-3', Reverse 5'-TCCCCATA¬CTCAGG¬CCTCTG-3'<br><br>*exon 2 - exon 4*: Forward 5'-GCCTCATGACCTCATGCCTG-3', Reverse 5'-AGCTTA¬GACGTAGGCACTGC-3'<br><br>*exon 5:* Forward 5'-GTCCCACAATAAGGGC¬TGCAC-3', Reverse 5'-CTGGGACAGAGGGCGAATGG-3' | This paper | NM_003018 |

| | | |
|---|---|---|
| SFTPC Genotyping, Primers for DNA Sequencing<br><br>*promoter + exon1:* 5'-CCCAGGTTTGCTCTTGCTGG-3'<br><br>*promoter + exon1*: 5'-GAGGAGGCAGGGCCCATCAC-3'<br><br>*exon 2*: 5'-TCCAGCCCTAGGACGCCGTG-3',<br><br>*exon 2*: 5'-CTGTCTGGCATGTCCTGTGC-3'<br><br>*exon 3:* 5'-GATGGGTACCACTGGCTGAG-3',<br><br>*exon 4*: 5'-TGGGTCAGGGAGAGAGCAGG-3',<br><br>*exon 4*: 5'-CACTCCTCCCAGCAGCCCTG-3'<br><br>*exon 5*: 5'-GTCCCACAATAAGGGCTGCAC-3'<br><br>*exon 5*: 5'-GGGAGTGGGAAGTACCGGTC-3'*exon 6*: 5'-CTGGGACAGAGGGCGAATGG-3' | This paper | NM_003018 |
